# Unravelling the role of thyroid hormones in seasonal neuroplasticity in European Starlings (Sturnus vulgaris)

**DOI:** 10.1101/2021.10.21.465109

**Authors:** Jasmien Orije, Sander R. Raymaekers, Gaurav Majumadar, Geert De Groof, Elisabeth Jonckers, Gregory F. Ball, Marleen Verhoye, Veerle M. Darras, Annemie Van der Linden

## Abstract

Thyroid hormones clearly play a role in the seasonal regulation of reproduction, but any role they might play in song behavior and the associated seasonal neuroplasticity in songbirds remains to be elucidated. To pursue this question, we first established seasonal patterns in the expression of thyroid hormone regulating genes in male European starlings employing *in situ* hybridization methods. Thyroid hormone transporter *LAT1* expression in the song nucleus HVC was elevated during the photosensitive phase, pointing towards an active role of thyroid hormones during this window of possible neuroplasticity. In contrast *DIO3* expression was high in HVC during the photostimulated phase, limiting the possible effect of thyroid hormones to maintain song stability during the breeding season. Next, we studied the effect of hypothyroidism on song behavior and neuroplasticity using *in vivo* MRI. Hypothyroidism inhibited the photostimulation-induced increase in testosterone, confirming the role of thyroid hormones in activating the hypothalamic–pituitary–gonadal (HPG) axis. Surprisingly, apart from the myelination of several tracts during the photostimulated phase, most neuroplasticity related to song production was unaffected by hypothyroidism. Remarkably, T3 plasma concentrations were negatively correlated to the microstructural changes in several song control nuclei. Potentially, a global reduction of circulating thyroid hormones during the photosensitive period is necessary to lift the brake imposed by the photorefractory period, whereas local fine-tuning of thyroid hormone concentrations through LAT1 could activate underlying neuroplasticity mechanisms. Given the complexity of thyroid hormone effects, this study is a steppingstone to disentangle the influence of thyroid hormones on seasonal neuroplasticity.

## 2 Introduction

Seasonal changes in steroid hormone concentrations such as testosterone play a central role in the regulation of song behavior and the associated neuroplasticity in seasonal songbirds (Balthazart, Charlier et al. 2010). For example, testosterone acting in the pre-optic area can increase the motivation to sing, resulting in activity-induced neuroplasticity, including increased forebrain song control nuclei volume (Alward, Balthazart et al. 2013) and stronger connections between the song control nuclei (Orije, Cardon et al. 2020). In several songbird species, testosterone is known to be necessary for the crystallization of song (Marler, Peters et al. 1988, Korsia and Bottjer 1991, Williams, Connor et al. 2003), while castration increases syllable’s entropy and decreases the song similarity and stereotypy (Wang, Liao et al. 2014). Male canaries sing a stable song during the breeding season, when testosterone levels are high. As canaries become photorefractory in fall, their testosterone levels decrease and their song becomes plastic again (Nottebohm, Nottebohm et al. 1987, Voigt and Leitner 2008, Alward, Mayes et al. 2014). Female canaries also produce song, when they are implanted with testosterone, their song rate increases and their song becomes stable. Female starlings implanted with testosterone show the same behavioral changes. Furthermore, when the testosterone implant is removed, song control nuclei rapidly decrease in volume (Orije, Cardon et al. 2020).

These studies demonstrate that testosterone can play a key role in stimulating songbirds of various species to sing at higher rates and to crystalize or stabilize their song, and that these behavioral changes are associated with structural changes in their song control system. The presence of aromatase, androgen and estrogen receptors in several song control nuclei (**Figure 10**) is consistent with the notion that testosterone can act directing in these key song control nuclei to induce changes in morphology and physiology. However, several studies have shown that even in absence or presences of very low concentrations of gonadal testosterone seasonal neuroplasticity can still occur, indicating that there are other factors that regulate or influence seasonal neuroplasticity (Smith, Brenowitz et al. 1997, Bentley, Van’t Hof et al. 1999, Dawson, King et al. 2001). One of the candidate factors that might regulate seasonal plasticity in brain and behavior are thyroid hormones. The avian thyroid gland produces thyroxine (T4) as well as a small amount of triiodothyronine (T3), which are both released into the bloodstream. Thyroid hormone activity is subsequently regulated at multiple levels. Several thyroid hormone transporters, including monocarboxylate transporters (MCT8 and MCT10), Na-independent organic anion transporting polypeptide 1C1 (OATP1C1), and L-type amino acid transporter 1 (LAT1) facilitate transport of T3 and T4 across the cell membrane (Heuer and Visser 2009). Once in the cell, the activity of thyroid hormones can be locally adjusted by deiodinases. Deiodinase 2 (DIO2) converts the prohormone T4 into its more bioactive form T3. Contrary, deiodinase 3 (DIO3) inactivates T4 and T3 to reverse T3 (rT3) and T2 respectively (Gereben, Zavacki et al. 2008). Thyroid hormones mostly operate by modulating gene expression as a result of binding of T3 to nuclear thyroid hormone receptors alpha or beta (THRA or THRB). These receptors associate with thyroid hormone response elements in the promotor region and induce or repress gene expression, depending on the gene.

Thyroid hormone levels change seasonally and are known to play a role in reproductive maturation and maintaining the photorefractory state in seasonally breeding birds (Bentley, Goldsmith et al. 1997, Bentley, Dawson et al. 2000). They closely interact with the hypothalamus-pituitary-gonadal (HPG) axis. One of the first things to change after exposure to a single long day of Japanese quail is the local upregulation of thyroid stimulating hormone (TSH) and DIO2 and downregulation of DIO3 in the pars tuberalis of the pituitary gland. This results in a local increase in T3 in the mediobasal hypothalamus, which causes gonadotropin releasing hormone (GnRH) release from morphologically changed GnRH nerve terminals and activates the rest of the HPG-axis (Yoshimura, Yasuo et al. 2003, Watanabe, Yamamura et al. 2007, Nakao, Ono et al. 2008). So far this link between thyroid hormones and gonadal steroid hormone production was not confirmed in starlings (Bentley, Tucker et al. 2013). Furthermore, thyroid hormones are known to play a role during neuronal development and are suggested to influence critical period learning (Bernal 2000, Yamaguchi, Aoki et al. 2012, Batista and Hensch 2019). However, the exact role of thyroid hormones in seasonal neuroplasticity in song and the song system has not been investigated yet.

We hypothesize that thyroid hormones contribute to the seasonal neuroplasticity related to song behavior. If thyroid hormones are indeed able to actively influence seasonal neuroplasticity, we expect in the first place expression of thyroid hormone receptors, transporters and deiodinases in the song bird brain and more specifically in the song control system. The first report describing a role of thyroid hormone regulating genes during development of the song control system and song learning in the zebra finch brain came from our coauthors (Raymaekers, Verbeure et al. 2017). They showed that DIO2 and LAT1 expression in several song control nuclei remained high during the sensory and sensorimotor phase of song learning in male zebra finches, suggesting that thyroid hormones are key players in this process. In line with the role of thyroid hormones in song development in zebra finches, we expected that the expression of thyroid hormone regulating genes could change seasonally in starlings and have differential effects in regulating song and neuroplasticity across the seasons. Once we had established the spatial and temporal expression patterns of thyroid hormone regulating genes (and thyroid hormone plasma levels) during different photoperiods, we induced hypothyroidism during the photosensitive stage to investigate the impact on seasonal neuroplasticity and song production (**Figure 1**).

**Figure 1:**
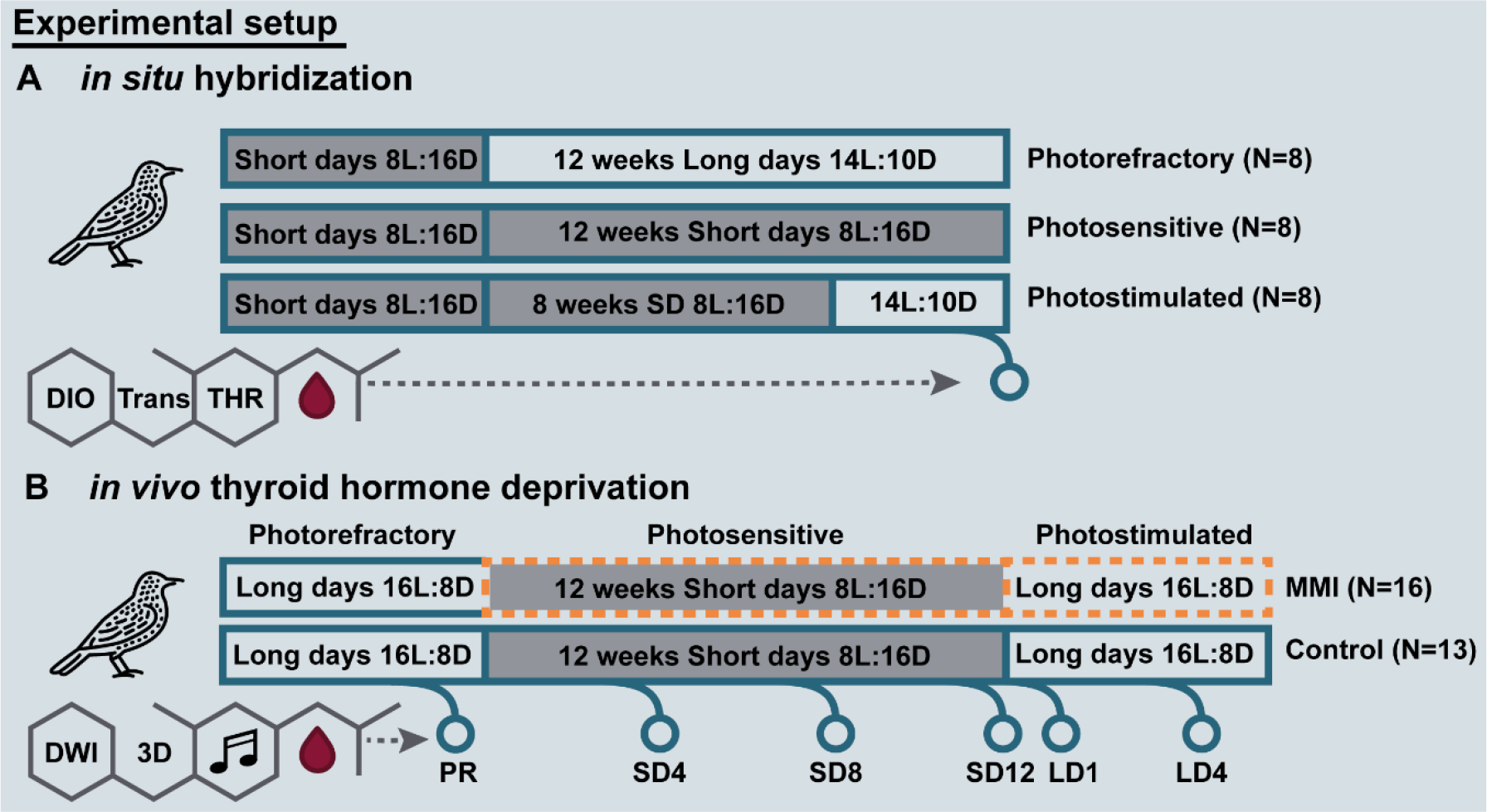
Overview experimental setup of in situ hybridization (A) and in vivo thyroid hormone deprivation **study.** For *in situ* hybridization, all birds were photosensitive after being on a short day photoperiod. These birds were divided into three groups with different photoperiodic manipulation: 12 weeks of long days to induce the photorefractory state, 12 weeks of short days to maintain photosensitive state, 8 weeks of short days followed by 4 weeks of long days to induce photostimulated state. For each photoperiod we analyzed different thyroid hormone regulators: deiodinases (DIO), thyroid hormone transporters (Trans), and thyroid hormone receptors (THR). For the in vivo manipulation study, we monitored individual starlings as they went through different photoperiods sequentially and diffusion weighted images (DWI) and 3D scans were acquired at the indicated time points. Short days (SD) are indicated by a grey box. Methimazole (MMI) treatment is indicated by an orange dotted line.

In short, this study aimed to answer the following questions: (1) Are thyroid hormone regulatory genes expressed in the song control system of male starlings? (2) Does their expression change seasonally? (3) Does thyroid hormone manipulation affect the HPG-axis just like in Japanese quail? (4) Does thyroid hormone deprivation affect seasonal neuroplasticity and song behavior in male starlings? We tried to answer the last question by inducing thyroid hormone deprivation at the onset of the photosensitive stage while using *in vivo* MRI to monitor neuroplasticity in starlings as they passed through different photoperiodic states. In a previous MRI study, we showed that seasonally increased neuroplasticity starts during the photosensitive period, which acts as a sensitive window for multisensory neuroplasticity as it shows involvement of the song control, visual and auditory system and even the cerebellum (Orije, Cardon et al. 2021). Furthermore, *in vivo* MRI is a powerful tool to study neural substrate changes associated with singing behavior or hormonal changes (Hamaide, Lukacova et al. 2020, Orije, Cardon et al. 2020, Orije, Cardon et al. 2021).

## 3 Results

### 3.1 Thyroid hormone regulators in the song control system: *in situ* hybridization

To understand fully the role and regulation of thyroid hormone action in seasonal neuroplasticity in male starlings, we performed ISH for mRNA of all known major thyroid hormone regulators. Of these thyroid hormone regulators *DIO2, MCT8, MCT10* and *OATP1C1* showed no visible expression in any of the photoperiods in any of the song nuclei nor in the rest of the telencephalon.

Of the main thyroid hormone activating and inactivating enzymes DIO2 and DIO3, only *DIO3* expression was detected in the HVC and varied over the different photoperiods (F(2, 18)=9.69 p=0.0014). Photorefractory and photosensitive birds showed little to no *DIO3+* cells, whereas *DIO3* expression was much higher after 4 weeks of photostimulation (**Figure 2A and Figure 3 A-C**).

**Figure 2:**
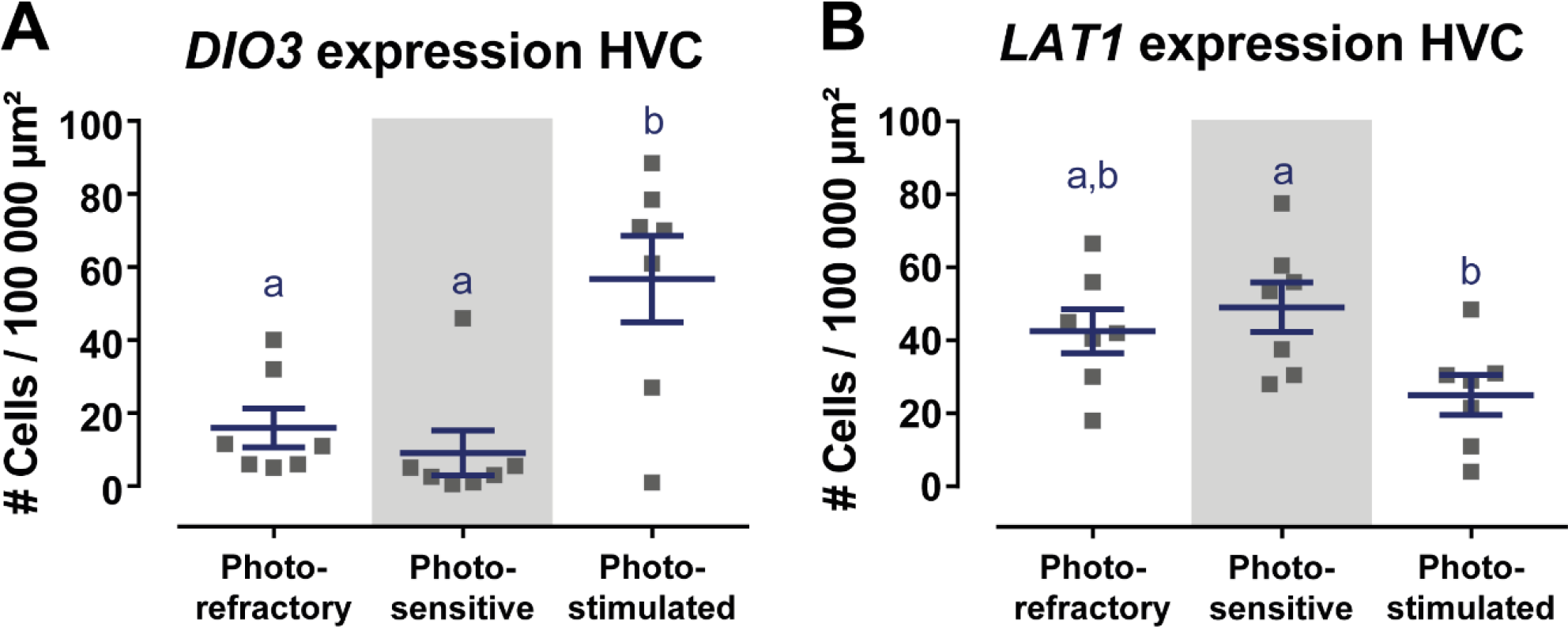
Overview of DIO3 (A) and LAT1 (B) expression in the HVC in the photorefractory, photosensitive and photostimulated phase. Dots represent the average number of DIO3+ and LAT1+ cells per individual bird. Horizontal bars represent the average with standard error of the mean error bars (n=7). The grey area indicates the photosensitive phase. Different letters denote significant differences by comparison with each other in post- hoc t-tests with p < 0.05 (multiple comparison correction with Dunn’s post hoc tests for DIO3 and Bonferroni correction for LAT1). If two time points share a common letter, DIO3 or LAT1 expression are not significantly different from each other.

**Figure 3:**
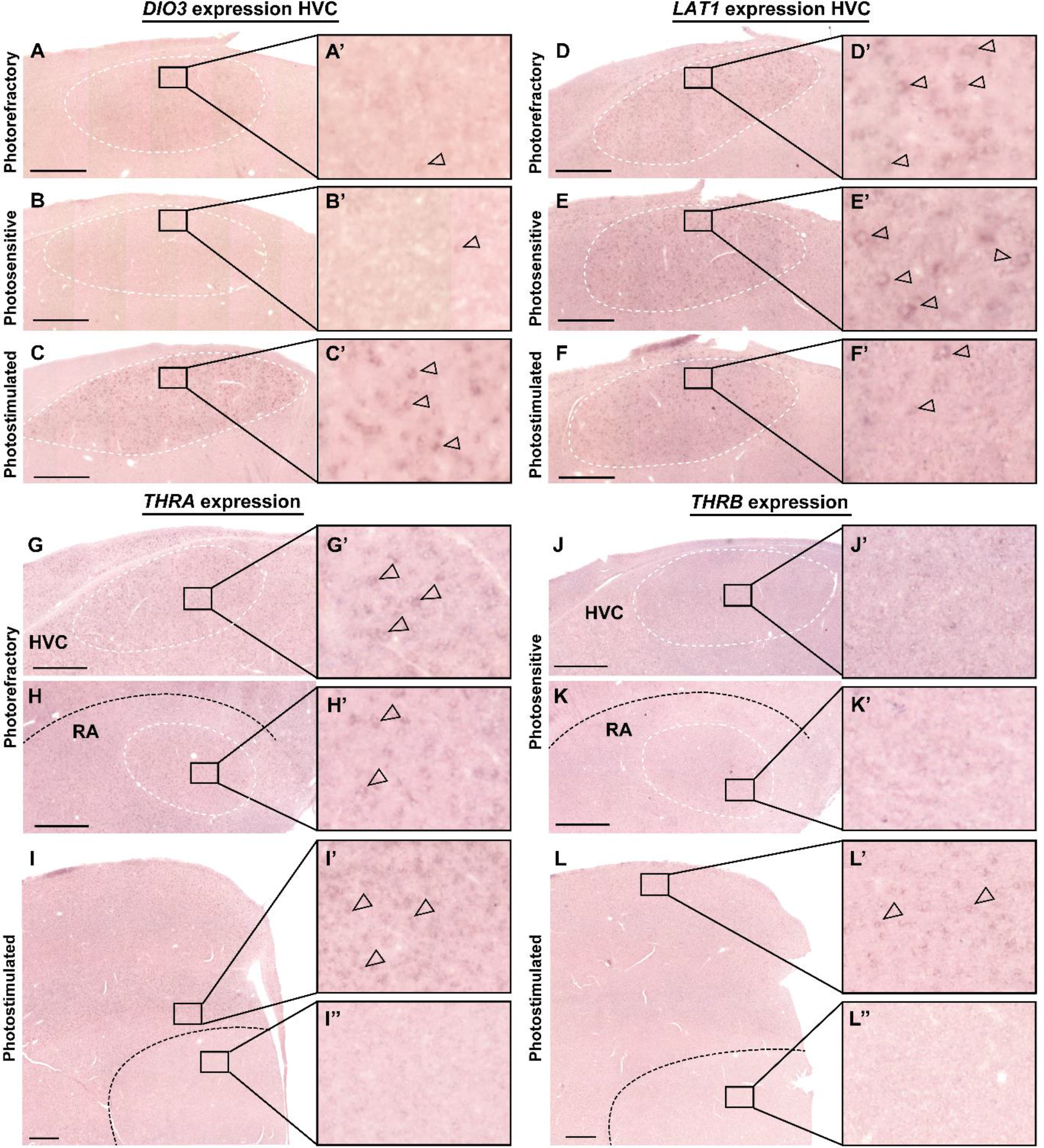
Representative images of *DIO3*. (A-C), *LAT 1* (D-F), *THRA* (G-I), *THRB* (J-L) mRNA expression in the starling brain. *DIO3* and *LAT1* mRNA expression changes seasonally over the photorefractory (A, D), photosensitive (B, E) and photostimulated phase (C, F). *THRA* and *THRB* mRNA expression remain unchanged during the different seasons in caudal telencephalon (G, H, J, K) at the level of HVC (G, J) and RA (H, K) and in the rostral telencephalon (I, L) at the level of LMAN (I’), pallium (L’) and Area X (I’’, L’’). The inserts indicate the location of the close-ups (A’-L’). The borders of HVC and RA are designated by white dashed lines. Black dashed line indicates the border between nidopallium and arcopallium in (H, K) and the border between striatum and pallium in (I, L). Empty arrowheads designate examples of DIO3+, LAT1+ and THRA+ cells respectively. Scale bar in overview pictures are 500µm.

In contrast to the other thyroid hormone transporters, *LAT1* exhibited a strong expression in HVC in all three photoperiods (F(2, 18)= 4.10, p=0.0342) (**Figure 3 D-F**). *Post hoc* Bonferroni test showed that the *LAT1* expression in the HVC in the photosensitive state was significantly higher than in the photostimulated phase (**Figure 2B**). Furthermore, there was a low, widespread expression of LAT1 in the upper layers of the pallium both rostrally and caudally and a low expression in the arcopallium including RA. No expression was observed in other song control nuclei.

Finally, we examined the expression pattern of thyroid hormone receptors. *THRA* showed a clear and widespread expression throughout the entire pallium including HVC, RA and LMAN, whereas no expression was detected in the striatum, including Area X (representative image shown in **Figure 3 G- I**). In general, there was no variation in *THRA* expression over the different photoperiods. Furthermore, the *THRA* expression was not specific for the song control nuclei, since similar density and intensity of *THRA* expression was found in the surroundings of HVC, RA and LMAN. *THRB* expression was much lower compared to *THRA* and was only found in upper layers of the pallium and caudal arcopallium but not detected in the RA or any other song control nucleus (**Figure 3 J-L**).

The widespread expression of *THRA* makes it possible for seasonal changes in T4 and T3 to affect several song control nuclei, except striatal regions like Area X. Active seasonal changes in thyroid hormone regulating genes (*DIO3* and *LAT1*) are confined to the HVC, as the central regulator of other song control nuclei it is connected to. This suggests that the intracellular concentration of thyroid hormones is actively controlled at the level of HVC, and could play a role in seasonal song behavior.

#### 3.1.1 Validation of ISH cell counting

To evaluate the validity of the cell count method used for the *DIO3* and *LAT1* ISH, 10 random sections of both ISHs were analyzed using the semi-quantitative manner of ‘stained surface fraction’. The correlation between both methods measuring expression (stained surface fraction values and original count value) was examined. The r² was 0.944 for the *DIO3* ISH and 0.924 for the *LAT1* ISH.

Furthermore, we investigated whether a change in cell density in the HVC across seasons could result in changes in *DIO3+* and *LAT1+* cell counts, by performing a Nissl staining on sections from each photoperiodic treatment in 3 birds. Cells stained by cresyl violet were counted in HVC, but numbers showed no variation over time (F(2, 20)=0.44, p=0.649) (Figure 3 – figure supplement 1).

### 3.2 Hypothyroidism by MMI treatment affects circulating thyroid hormones (T3 and T4) and testosterone concentrations

The aim of the MMI treatment was to inhibit thyroid hormone production in order to establish how hypothyroidism affects seasonal neuroplasticity. To confirm the effectiveness of this treatment, we took blood samples at each time point to monitor plasma levels of T3 and T4. At the baseline photorefractory time point there was no difference in circulating thyroid hormone concentrations between groups. MMI treatment was successful in reducing both T3 and T4 to a minimum at 4 weeks after starting MMI treatment upon the switch to short days (**Figure 4 A, B**). Thyroid hormone levels remained low for the rest of the experiment. In the control group T4 and T3 significantly decreased during the photosensitive period compared to the photorefractory period. After switching back to long days, T4 concentrations significantly increased again to levels comparable to the photorefractory period, while T3 concentrations remained low.

**Figure 4:**
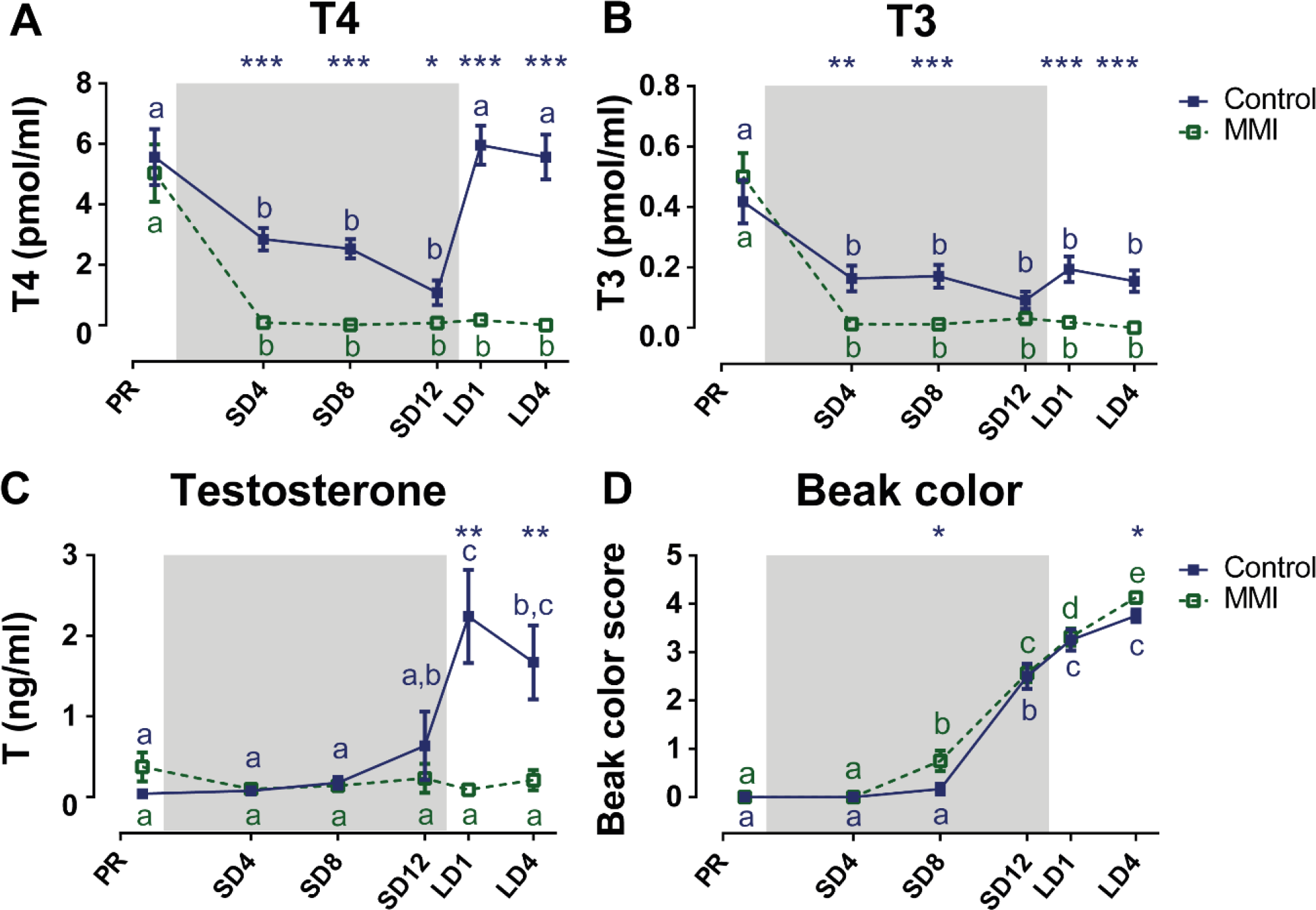
Overview of the seasonal changes in plasma levels of thyroid hormones T4 (A), T3 (B), testosterone and beak color (D) in control and MMI-treated starlings. Solid and dashed lines represent the group average of control and MMI-treated starlings respectively with standard errors of the mean error bars. The grey area indicates the photosensitive period of short days (8L:16D). Different letters denote significant differences by comparison with each other in post-hoc t-tests with p < 0.05 (Tukey’s HSD correction for multiple comparisons) comparing the different time points to each other. If two time points share a common letter, the thyroid hormone or testosterone levels are not significantly different from each other.

Additionally, we monitored circulating testosterone concentrations to determine how photoperiod and MMI treatment affect the HPG-axis. While the HPG-axis matured in the control group as shown by the elevated circulating testosterone levels during the photostimulated phase, circulating testosterone level remained low throughout the different photoperiods in the MMI-treated group (**Figure 4 C**). However, the beak color of MMI-treated birds turned yellow at the end of the photosensitive phase and in the photostimulated phase just like in the control group (**Figure 4 D**). Beak color does not directly represent the amount of circulating testosterone, but rather indicates that testosterone is present in circulation, even when in low doses below the sensitivity limit of the RIA assay (Ball and Wingfield 1987, Wingfield and Silverin 2002). Overall, MMI treatment does not only decrease thyroid hormone but also testosterone plasma concentrations. However, testosterone was not totally eliminated, as can happen with castration, as indicated by the beak color changing to yellow in the MMI treated birds. Overall these data indicate that thyroid hormones indeed influence HPG-axis physiology in starlings as hypothesized previously based on work in other species (Yoshimura 2013).

### 3.3 Hypothyroidism by MMI treatment differentially affects song rate and song bout length

We also investigated how MMI treatment affects the change in song behavior over different photoperiods. Both control and MMI-treated male starlings sang a longer song bout during the photostimulated phase compared to the photosensitive phase, but MMI-treated starlings further increased their song bout length during the photostimulated phase from 35.26±0.62 seconds at LD1 to 39.36±0.71 seconds at LD4 (**Figure 5**). Furthermore, song bout length was within-subjects positively correlated with the testosterone concentration in the control group (rmcorr=0.572, p=0.0054). Since testosterone plasma concentration in the MMI-treated group did not change over different seasons, it was not correlated to song bout length. However, beak color, which is a sensitive indicator of the presence or absence of circulating testosterone, did positively correlate to song bout length in both groups (control: rmcorr=0.592, p=0.0015; MMI: rmcorr=0.4136, p=0.0287). Unlike the song bout length, the song rate did not change over time nor did it correlate to hormone levels in either group. However, the untreated male starlings had a higher song rate compared to MMI-treated starlings after 4 weeks of photostimulation.

**Figure 5:**
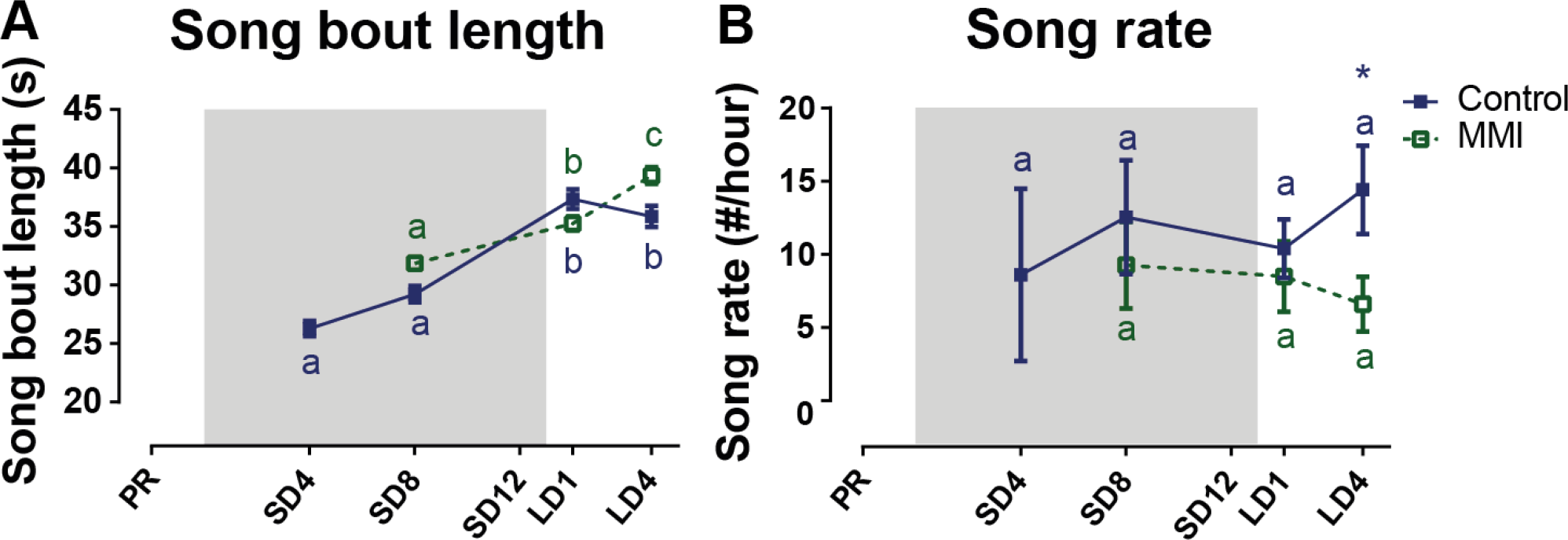
Overview of the seasonal changes in song bout length (A) and song rate (B) in control and MMI-treated starlings. Solid and dashed lines represent the group average of control and MMI-treated starlings respectively with standard errors of the mean error bars. The grey area indicates the photosensitive period of short days (8L:16D). Different letters denote significant differences by comparison with each other in post-hoc t-tests with p < 0.05 (Tukey’s HSD correction for multiple comparisons) comparing the different time points to each other. If two time points share a common letter, the song bout length and song rate are not significantly different from each other.

### 3.4 Effect of hypothyroidism on seasonal neuroplasticity: a longitudinal DTI study

In order to determine whether thyroid hormone deprivation affects neuroplasticity, we used *in vivo* DWI to monitor structural neuroplasticity changes within the starling brain, using the same method as Orije, Cardon et al. (2021). This method measures several diffusion parameters that can provide information on microstructural changes related to structure and spatial organization of neuronal cells and myelin (Orije, Cardon et al. 2020, Orije, Cardon et al. 2021). A voxel based factorial analysis was performed to establish the general fractional anisotropy changes over time and if this differed between both experimental groups, it was indicated by a significant interaction. The results are summarized in **Figure 6, Figure 7 and Table 1.** From each significant cluster, we extracted the mean fractional anisotropy values and plotted them over time to determine the profile of temporal changes in fractional anisotropy (**Figure 6, Figure 7 B, D**). Additionally, we extracted other diffusion parameters (mean, axial and radial diffusivity) and fixel-based measures from these ROIs to provide more insight into the basis of the fractional anisotropy change (**Figure 6 - figure supplements 1-2, Figure 7 - figure supplements 1-2**). For the statistical analysis of the extracted parameters, we used linear mixed model analysis with Tukey’s Honest Significant Difference (HSD) as multiple comparison correction during *post hoc* statistics.

**Figure 6:**
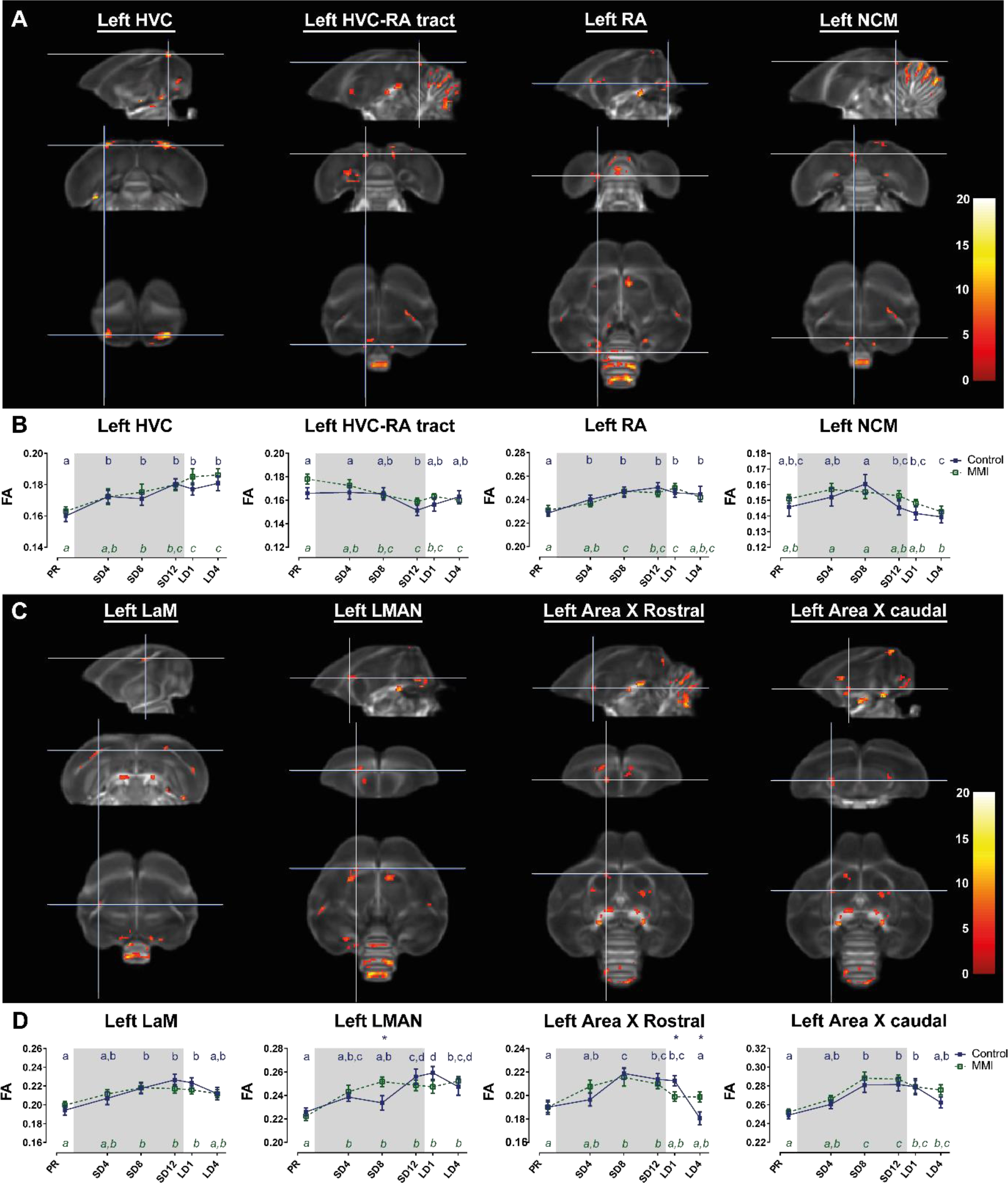
Voxel-based flexible factorial analysis revealed time effect in the song motor pathway and auditory pathway (A), and anterior forebrain pathway (C). The longitudinal changes over time of the extracted diffusion parameter fractional anisotropy of each ROI-based cluster is displayed below (B, D). The statistical maps were assessed at puncorr<0.001 and kE≥20 voxels with a small volume correction including regions of the song control system, white matter structures and the cerebellum. The grey area indicates the photosensitive period of short days (8L:16D). *Post hoc* statistical testing with Tukey’s HSD multiple comparison (p < 0.05) correction revealed significant differences between different time points, visualized by different letters. If two time points share the same letter, the DTI values are not significantly different from each other. Significant group differences at specific time points are indicated by *(p < 0.05), ** (p < 0.01) or *** (p < 0.001). Error bars shown are the standard error of the mean.

**Figure 7:**
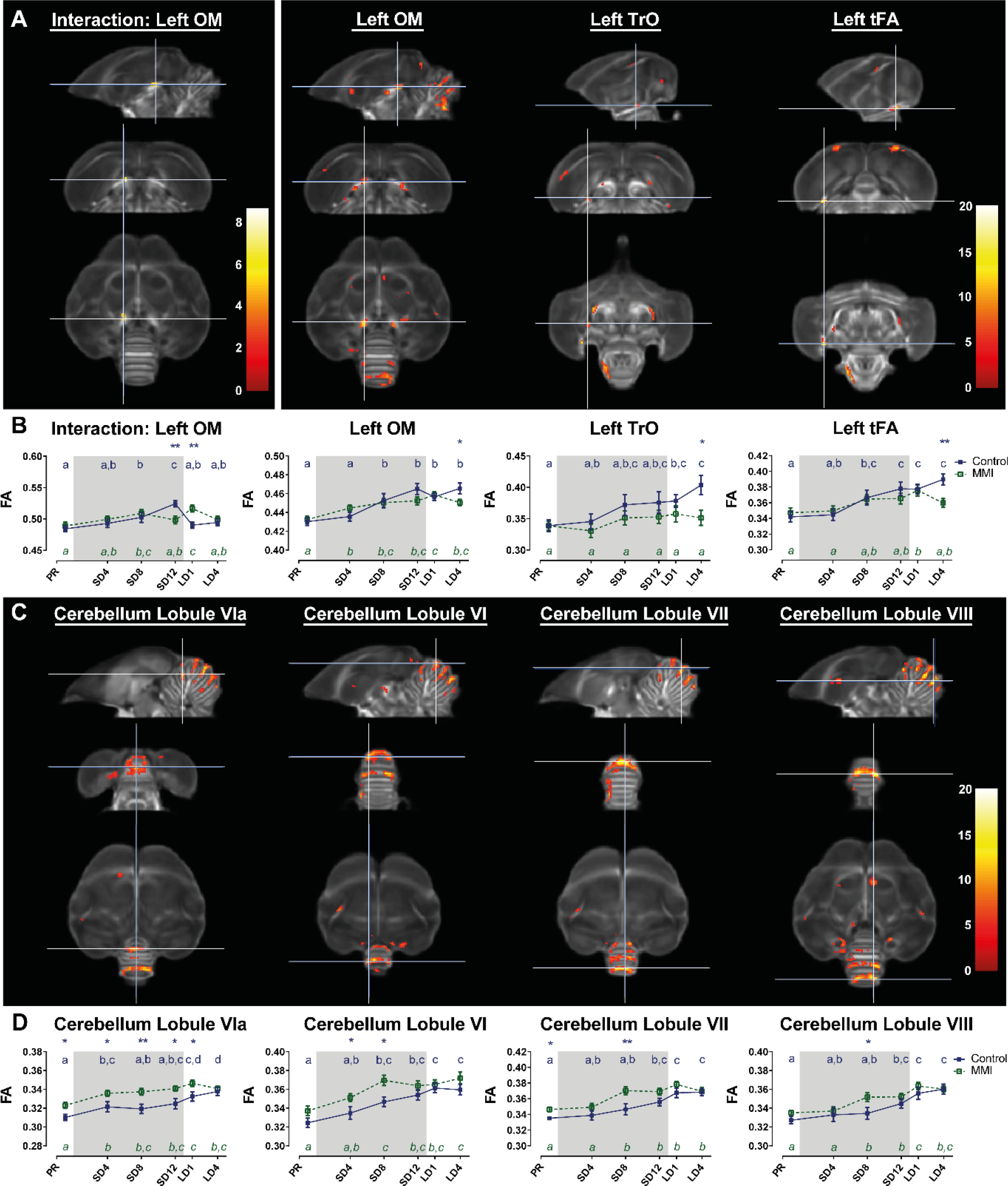
Voxel-based flexible factorial analysis revealed an interaction between time and treatment group in a specific part of the OM tract and time effect in the white matter tracts (A), and specific cerebellar lobules (C). The longitudinal changes over time of the extracted diffusion parameter fractional anisotropy of each ROI-based cluster is displayed below (B, D). The statistical maps were assessed at puncorr<0.001 and kE≥20 voxels with a small volume correction including regions of the song control system, white matter structures and the cerebellum. The grey area indicates the photosensitive period of short days (8L:16D). *Post hoc* statistical testing with Tukey’s HSD multiple comparison (p < 0.05) correction revealed significant differences between different time points, visualized by different letters. If two time points share the same letter, the DTI values are not significantly different from each other. Significant group differences at specific time points are indicated by *(p < 0.05), ** (p < 0.01) or *** (p < 0.001). Error bars shown are the standard error of the mean.

**Table 1:** Overview interaction and time effect in fractional anisotropy in small volume correction.

| Effect | Cluster | Hemisphere | Cluster |  | Peak |  |
| --- | --- | --- | --- | --- | --- | --- |
| | | | $p_{FWE}$ | $k_E$ | $p_{FWE}$ | $F$ |
| Interaction time * group | OM | Left | 0.066 | 25 | 0.017 | 8.71 |
| Main time effect | HVC surroundings | Left | <0.0001 | 80 | <0.0001 | 14.28 |
|  |  | Right | <0.0001 | 233 | <0.0001 | 16.71 |
|  | OM | Left | <0.0001 | 196 | <0.0001 | 13.45 |
|  |  | Left | 0.001 | 63 | 0.007 | 9.18 |
|  |  | Right | <0.0001 | 124 | 0.002 | 9.98 |
|  |  | Right | 0.032 | 30 | 0.003 | 9.72 |
|  | Area X rostral surroundings | Left | 0.007 | 41 | 0.266 | 7.12 |
|  |  | Right | <0.0001 | 123 | <0.0001 | 10.86 |
|  | Area X caudal surroundings | Left | 0.002 | 52 | 0.054 | 8.04 |
|  |  | Right | <0.0001 | 94 | 0.533 | 6.55 |
|  | tFA | Left | 0.001 | 57 | <0.0001 | 16.39 |
|  |  | Right | 0.003 | 49 | 0.001 | 10.14 |
|  | Optic tract | Left | 0.004 | 45 | <0.0001 | 17.02 |
|  |  | Left | 0.037 | 29 | 0.003 | 9.74 |
|  |  | Right | 0.003 | 48 | 0.001 | 10.39 |
|  | RA surroundings | Left | <0.0001 | 96 | 0.014 | 8.80 |
|  |  | Left | 0.008 | 40 | 0.038 | 8.25 |
|  |  | Left | 0.028 | 31 | 0.025 | 8.48 |
|  |  | Right | 0.077 | 24 | 0.043 | 8.18 |
|  | HVC-RA tract | Left | 0.003 | 49 | 0.053 | 8.06 |
|  |  | Right | 0.002 | 52 | 0.021 | 8.59 |
|  | LMAN | Left | <0.0001 | 157 | 0.026 | 8.45 |
|  | Lamina mesopallialis | Left | 0.003 | 49 | 0.024 | 8.49 |
|  |  | Left | 0.001 | 57 | 0.132 | 7.54 |
|  |  | Right | 0.057 | 26 | 0.039 | 8.23 |
|  | Caudal NCM | Left | 0.011 | 38 | 0.225 | 7.24 |
|  | Cerebellum | Lobule VIII | <0.0001 | 528 | <0.0001 | 20.20 |
|  |  | Lobule VII | <0.0001 | 741 | <0.0001 | 19.92 |
|  |  | Lobule VIa | <0.0001 | 271 | <0.0001 | 13.21 |
|  |  | Left side | <0.0001 | 255 | <0.0001 | 13.15 |
|  |  | Lobule VI | <0.0001 | 419 | <0.0001 | 12.46 |

Table summarizing the significance at cluster level and peak level. P-values are FWE corrected. $k_E$ indicates the number of voxels within a cluster. Grey values indicate clusters that are significant at cluster level but not at peak level.

Overall most neuroplasticity changes looked similar in both groups. In line with the prior longitudinal MRI study (Orije, Cardon et al. 2021), the fractional anisotropy value of several parts of the song motor pathway, anterior forebrain pathway but also visual system and molecular layer of the cerebellum increase gradually over time, starting during the photosensitive period. The MMI-treated group had higher FA values in the molecular layer of the cerebellum at several time points, including the first time point. Since the difference was already present at the start of the experiment, before the start of MMI treatment, it most probably reflects unintended sample variation between groups. Despite this unintended group difference, both groups experienced a similar increase in FA over time.

Only a small part of the left OM showed a significant interaction between time and group. The control group peaked in fractional anisotropy at SD12, whereas the MMI-treated group reached a maximum at LD1. *Post hoc* linear mixed model analysis revealed more subtle interactions over time in the different groups. In the rostral part of the left Area X surroundings, the control group had a decrease in fractional anisotropy at LD4, whereas in the MMI group fractional anisotropy remained constant. In the MMI group fractional anisotropy values in LMAN significantly increased at SD4, whereas fractional anisotropy values in the control group only increased at the end of the photosensitive period. Most interestingly, several tracts including the OM, tFA and optic tract have similar *post hoc* interactions, where the control group increased in fractional anisotropy at LD4, but the MMI-treated group did not. This increase in fractional anisotropy was mainly linked to a decrease in radial diffusivity at LD4. At LD4 there was a significant difference between both groups in fractional anisotropy and radial diffusivity. This indicates that the increase in fractional anisotropy at LD4 in the control group was most probably due to increased myelination of these tracts, something that is missing in the MMI- treated group.

In conclusion, MMI-induced hypothyroidism did not prevent the overall neuroplasticity to occur during the photosensitive period. Only during the photostimulated phase myelination of several tracts was affected in the MMI-treated group.

### 3.5 Correlations between song behavior, its neural substrate and thyroid hormones

After describing the seasonal neuroplasticity in a general way, we aimed at determining how these neuroplastic fractional anisotropy changes relate to changes in song behavior and hormone levels. Voxel-based multiple regression between song behavior and fractional anisotropy values in the brain was used to determine the location of neural correlates. Interestingly, the song bout length showed a positive correlation to fractional anisotropy values at the level of the left HVC, RA surroundings (bilaterally), left LMAN, left part of the lateral septum (LS) and cerebellar lobules VI-VII whereas a cluster near the HVC-RA tract was negatively correlated to the song bout length (**Figure 8, Table 2**). After extraction of the fractional anisotropy values of the significant clusters, this information was used to determine the within and between-subject correlation of each group separately. This way, we tried to distinguish whether the correlation was present in both groups. Both groups had significant between-subject correlations in all regions, except for the LS that only had a significant between-subject and within-subject correlation in the control group but not in the MMI-treated group (**Table 3**). The correlation to song bout length was largely driven by a between-subject correlation, meaning that birds with longer song bout lengths, had higher fractional anisotropy values in song control nuclei, cerebellar lobules VI-VII and LS. Furthermore, the fractional anisotropy values of the right RA, LMAN and cerebellar lobules VI-VII were also correlated on a within-subject level in both the control and MMI-treated group. Additionally, MMI-treated starlings present a repeated measures correlation at the level of Left RA and right HVC-RA tract, indicating that the variation over time in song bout length was also correlated to the variation in structural neuroplasticity (measured by fractional anisotropy) on a subject level.

**Figure 8:**
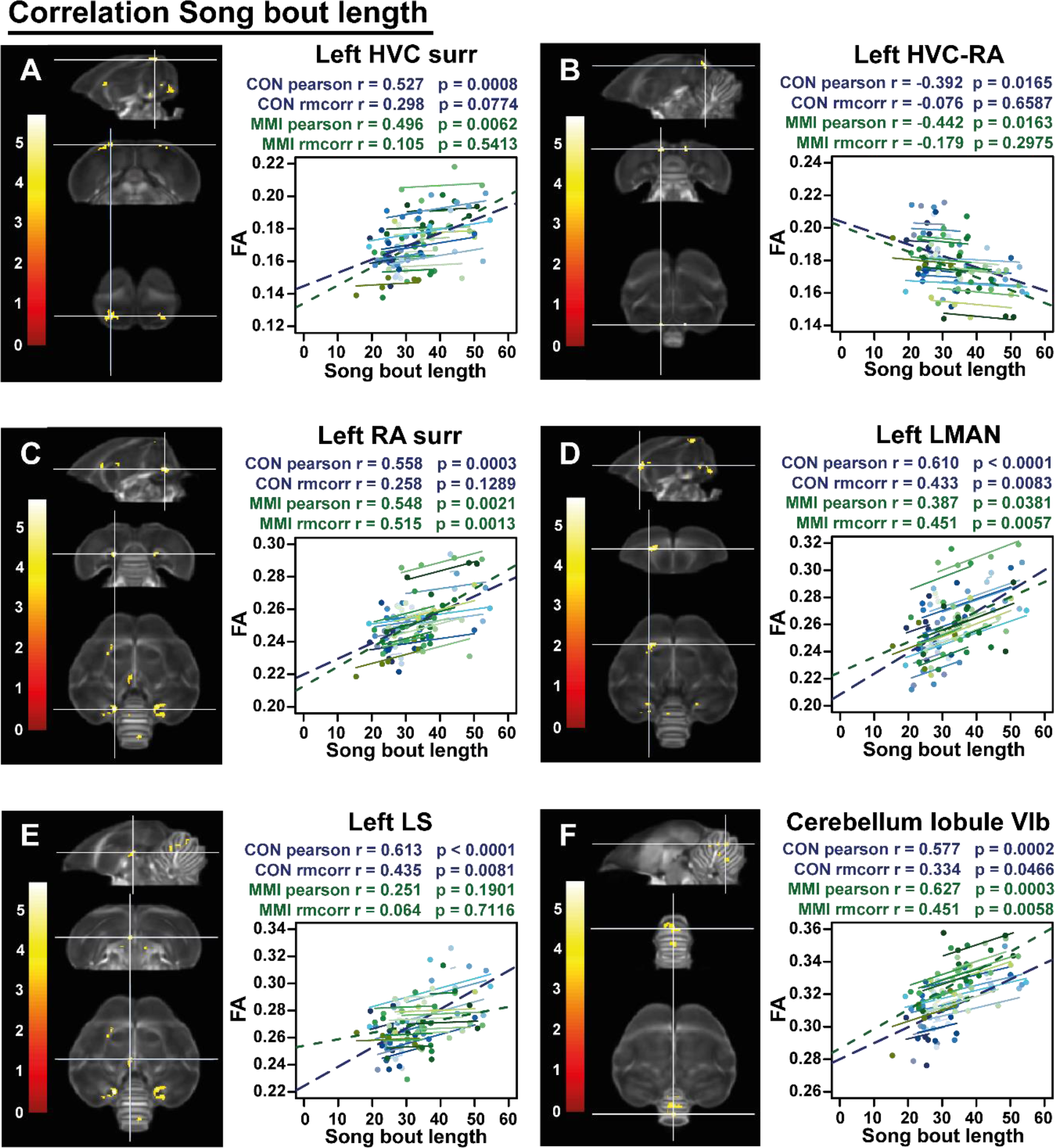
Overview of structural neural correlates of song bout length to fractional anisotropy in both groups identified using voxel-based multiple regression. The statistical maps were assessed at puncorr<0.001 and kE≥20 voxels with a small volume correction including regions of the song control system, white matter structures and the cerebellum. Below each statistical parametric map, the identified correlations were further explored with repeated measures correlation of the control group (blue) and MMI treatment group (green). Solid colored lines show the best linear fit for the within-subject correlation in the control group (blue) and MMI group (green) using parallel regression lines for individual animals. The blue and green dashed line represent the linear fit of the overall Pearson correlation representing the between-subject correlations of the control and MMI group respectively. For bilateral correlations, only the left side is shown. Abbreviations: surr, surroundings.

**Table 2:** Summary of the voxel-based correlation analysis of song bout length and thyroid hormones T3 and T4.

| Effect | Cluster | Hemisphere | Cluster |  | Peak |  |
| --- | --- | --- | --- | --- | --- | --- |
| | | | $p_{FWE}$ | $k_E$ | $p_{FWE}$ | $T$ |
| Song bout length | RA surr | Left | <0.0001 | 192 | 0.005 | 5.68 |
|  |  | Right | <0.0001 | 220 | 0.016 | 5.39 |
|  | HVC surr | Left | <0.0001 | 136 | 0.006 | 5.63 |
|  | LMAN | Left | <0.0001 | 134 | 0.022 | 5.31 |
|  | Cerebellum | Lobule VIa | 0.005 | 79 | 0.224 | 4.58 |
|  |  | Lobule VIb | <0.0001 | 157 | 0.069 | 4.97 |
|  |  | Lobule VII | 0.012 | 65 | 0.074 | 4.95 |
|  |  | Lobule VIII | 0.045 | 47 | 0.190 | 4.64 |
|  | LS | Left | 0.033 | 51 | 0.472 | 4.29 |
|  | HVC-RA-tract | Left | 0.036 | 50 | 0.081 | 4.92 |
|  |  | Right | 0.161 | 31 | 0.023 | 5.30 |
| T3 | HVC surr | Right | 0.005 | 136 | 0.003 | 5.54 |
|  | Area X Csur | Right | 0.004 | 84 | 0.009 | 5.28 |
|  | RA surr | Left | 0.227 | 28 | 0.047 | 4.86 |
|  | Cerebellum | Lobule VIb | <0.0001 | 209 | 0.008 | 5.31 |
|  |  | Lobule VI | <0.0001 | 132 | 0.011 | 5.22 |
|  |  | Lobule VIa | <0.0001 | 171 | 0.062 | 4.78 |
|  |  | Lobule VII | 0.001 | 104 | 0.113 | 4.62 |
| T4 | HVC surr | Right | 0.056 | 46 | 0.012 | 5.24 |
|  | Cerebellum | Lobule V | <0.0001 | 214 | 0.098 | 4.68 |

This table summarizes for each significant cluster-based ROI the voxel-based FWE corrected p-value at cluster and peak level, next to the overall Pearson's correlation, the repeated measures correlation (rmcorr) and its significance. $k_E$ indicates the cluster size. $T$ reflects the t-value of the maximal peak located within the cluster-based ROI. Voxel-based correlations in grey are significant at the cluster level, but not at peak level. Abbreviations: C, caudal; surr, surroundings.

**Table 3:**
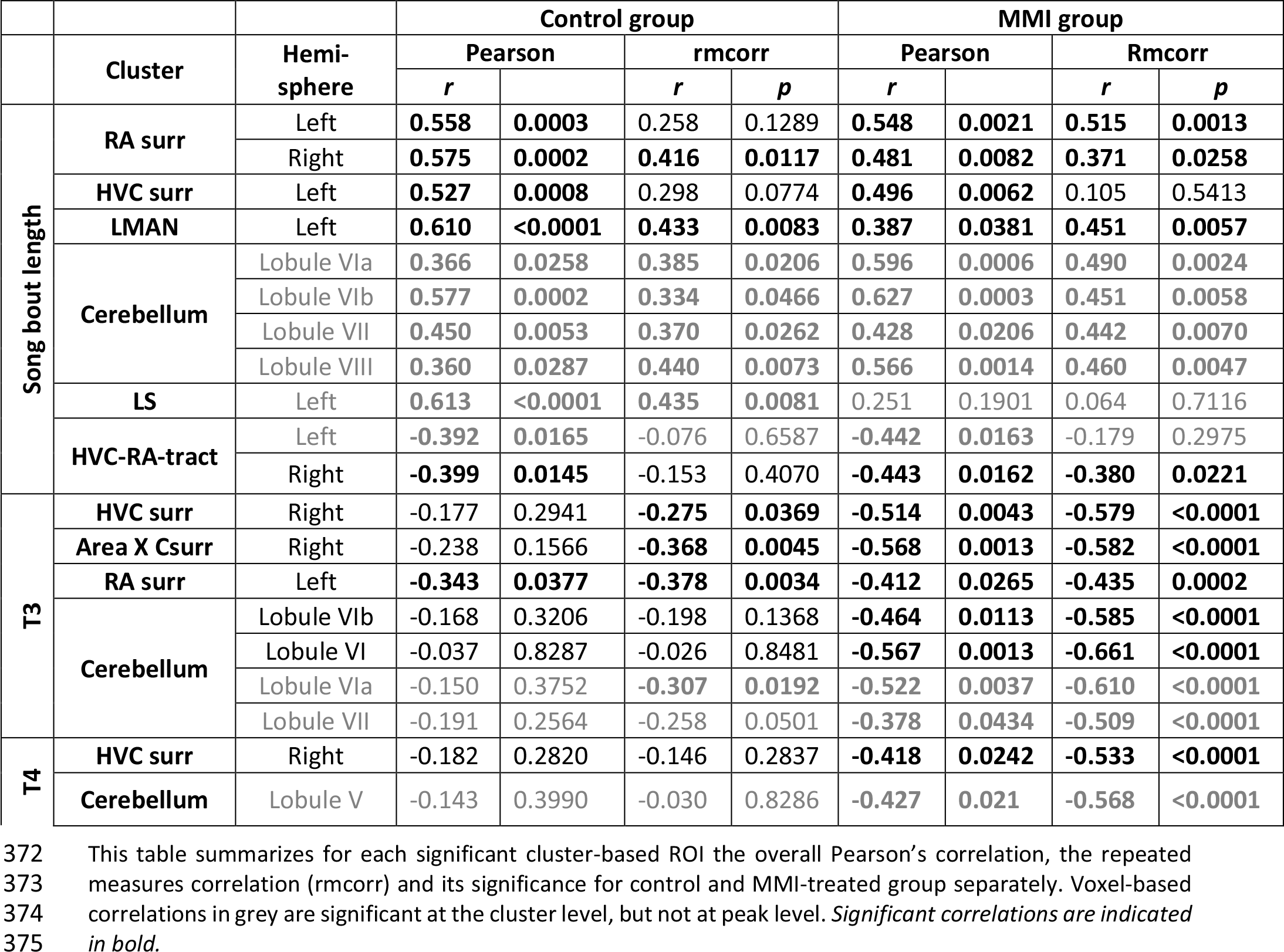
Summary of the overall and repeated measures correlation of song bout length and thyroid hormones T3 and T4 for control and MMI-treated group separately.

|  | Cluster | Hemi-<br>sphere | Control group |  |  |  | MMI group |  |  |  |
| --- | --- | --- | --- | --- | --- | --- | --- | --- | --- | --- |
|  |  |  | Pearson |  | rmcorr |  | Pearson |  | Rmcorr |  |
|  |  |  | *r* |  | *r* | *p* | *r* |  | *r* | *p* |
| Song bout length | RA surr | Left | **0.558** | **0.0003** | 0.258 | 0.1289 | **0.548** | **0.0021** | **0.515** | **0.0013** |
|  |  | Right | **0.575** | **0.0002** | **0.416** | **0.0117** | **0.481** | **0.0082** | **0.371** | **0.0258** |
|  | HVC surr | Left | **0.527** | **0.0008** | 0.298 | 0.0774 | **0.496** | **0.0062** | 0.105 | 0.5413 |
|  | LMAN | Left | **0.610** | **<0.0001** | **0.433** | **0.0083** | **0.387** | **0.0381** | **0.451** | **0.0057** |
|  | Cerebellum | Lobule VIa | **0.366** | **0.0258** | **0.385** | **0.0206** | **0.596** | **0.0006** | **0.490** | **0.0024** |
|  |  | Lobule VIb | **0.577** | **0.0002** | **0.334** | **0.0466** | **0.627** | **0.0003** | **0.451** | **0.0058** |
|  |  | Lobule VII | **0.450** | **0.0053** | **0.370** | **0.0262** | **0.428** | **0.0206** | **0.442** | **0.0070** |
|  |  | Lobule VIII | **0.360** | **0.0287** | **0.440** | **0.0073** | **0.566** | **0.0014** | **0.460** | **0.0047** |
|  | LS | Left | **0.613** | **<0.0001** | **0.435** | **0.0081** | 0.251 | 0.1901 | 0.064 | 0.7116 |
|  | HVC-RA-tract | Left | **-0.392** | **0.0165** | -0.076 | 0.6587 | **-0.442** | **0.0163** | -0.179 | 0.2975 |
|  |  | Right | **-0.399** | **0.0145** | -0.153 | 0.4070 | **-0.443** | **0.0162** | **-0.380** | **0.0221** |
| T3 | HVC surr | Right | -0.177 | 0.2941 | **-0.275** | **0.0369** | **-0.514** | **0.0043** | **-0.579** | **<0.0001** |
|  | Area X Csur | Right | -0.238 | 0.1566 | **-0.368** | **0.0045** | **-0.568** | **0.0013** | **-0.582** | **<0.0001** |
|  | RA surr | Left | **-0.343** | **0.0377** | **-0.378** | **0.0034** | **-0.412** | **0.0265** | **-0.435** | **0.0002** |
|  | Cerebellum | Lobule VIb | -0.168 | 0.3206 | -0.198 | 0.1368 | **-0.464** | **0.0113** | **-0.585** | **<0.0001** |
|  |  | Lobule VI | -0.037 | 0.8287 | -0.026 | 0.8481 | **-0.567** | **0.0013** | **-0.661** | **<0.0001** |
|  |  | Lobule VIa | -0.150 | 0.3752 | **-0.307** | **0.0192** | **-0.522** | **0.0037** | **-0.610** | **<0.0001** |
|  |  | Lobule VII | -0.191 | 0.2564 | -0.258 | 0.0501 | **-0.378** | **0.0434** | **-0.509** | **<0.0001** |
| T4 | HVC surr | Right | -0.182 | 0.2820 | -0.146 | 0.2837 | **-0.418** | **0.0242** | **-0.533** | **<0.0001** |
|  | Cerebellum | Lobule V | -0.143 | 0.3990 | -0.030 | 0.8286 | **-0.427** | **0.021** | **-0.568** | **<0.0001** |

This table summarizes for each significant cluster-based ROI the overall Pearson's correlation, the repeated measures correlation (rmcorr) and its significance for control and MMI-treated group separately. Voxel-based correlations in grey are significant at the cluster level, but not at peak level. *Significant correlations are indicated* *in bold.*

Remarkably many of the neuroplasticity changes in the surroundings of the song control nuclei and cerebellum were negatively correlated to the levels of circulating T3 and T4 (**Figure 9, Table 2**). Right HVC, left RA, right Area X surroundings and cerebellar lobules VI-VII were negatively correlated to the circulating T3 levels in both the MMI and control group. As circulating T3 levels decreased during the photosensitive period, whether naturally in the control group or artificially in the MMI-treated group, fractional anisotropy values increased in these regions. This correlation was mostly driven by a between and within-subject correlation in the MMI-treated group (**Table 3**). Only the RA had a significant negative between and within-subject correlation in the control group. Additionally, the control group had significant within-subject correlations to T3 at the level of the right HVC and caudal part of Area X surroundings. Correlations between T4 and the FA values at HVC surroundings and cerebellum only existed in the MMI-treated group. Rather than the natural biological variation in T4 in the control group, it is the artificial lowering of the T4 levels by the MMI treatment that correlated to the neuroplasticity in these regions.

**Figure 9:**
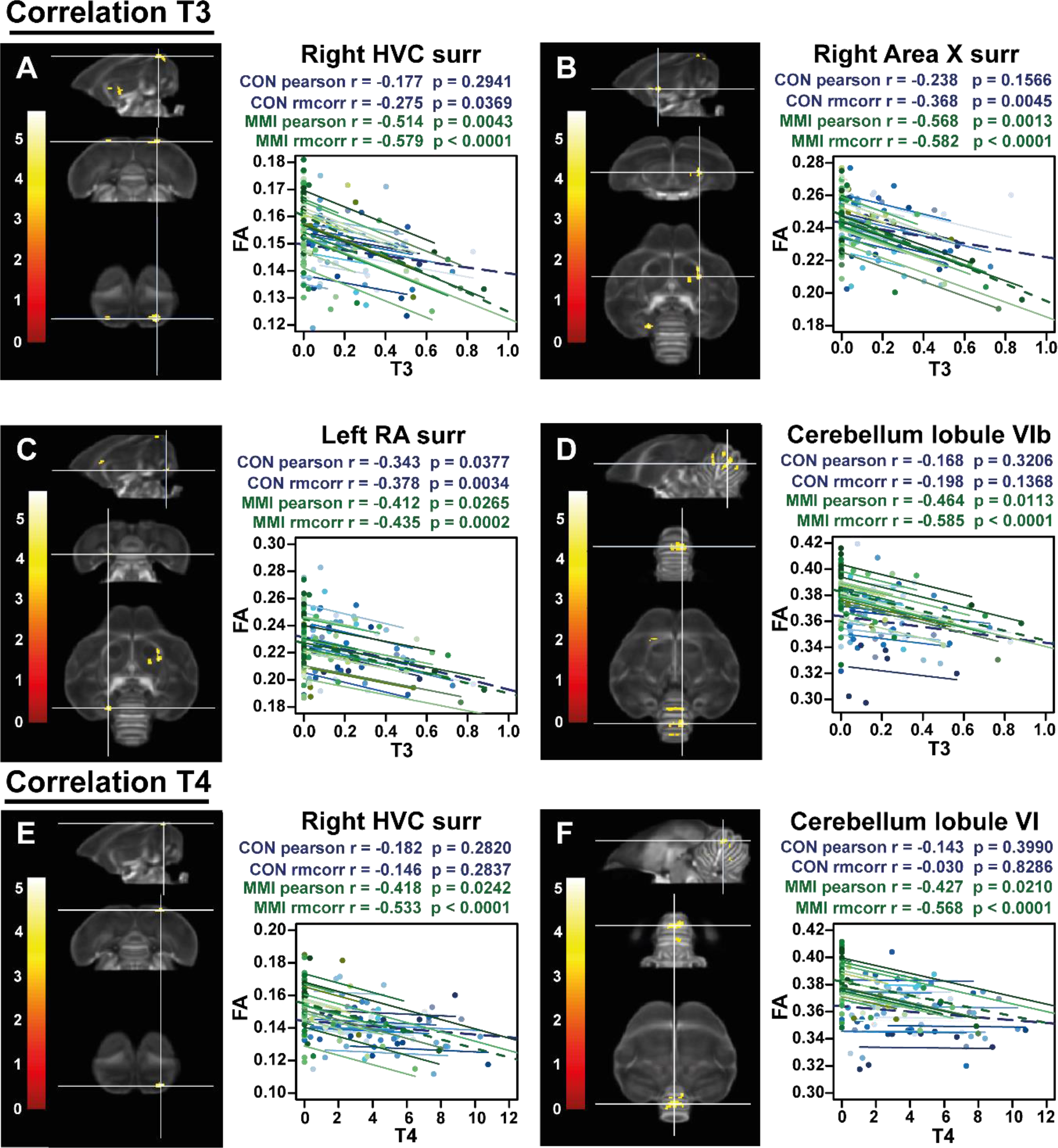
Overview of structural neural correlates of thyroid hormones T3 (A-D) and T4 (E-F) levels to fractional anisotropy in both groups identified using voxel-based multiple regression. The statistical maps were assessed at puncorr<0.001 and kE≥20 voxels with a small volume correction including regions of the song control system, white matter structures and the cerebellum. Next to each statistical parametric map, the identified correlations were further explored with repeated measures correlation of the control group (blue) and MMI treatment group (green). Solid colored lines show the best linear fit for the within-subject correlation in the control group (blue) and MMI group (green) using parallel regression lines for individual animals. The blue and green dashed line represent the linear fit of the overall Pearson correlation representing the between-subject correlations of the control and MMI group respectively. For bilateral correlations, only the left side is shown. Abbreviations: surr, surroundings.

## 4 Discussion

We demonstrate in this study that there are indeed thyroid hormone regulating genes expressed in the song control system, and that expression changes seasonally in the HVC. High *LAT1* expression during the photosensitive phase points to an increased uptake of thyroid hormones in the HVC during this sensitive window of neuroplasticity, whereas during photostimulation, elevated *DIO3* expression limits the amount of active thyroid hormone, presumably to sustain song stability during the breeding season. To establish further the effect of thyroid hormones on song behavior and neuroplasticity, we induced hypothyroidism by MMI treatment. As expected, hypothyroidism inhibited the photostimulation-induced increase in testosterone concentrations in the blood, indicating that as in Japanese quail thyroid hormones play a role in the activation of the HPG-axis upon photostimulation in starlings. The effect of thyroid hormones on testosterone is important for the interpretation of song behavioral changes, since testosterone is known for its effect on song motivation and song quality measures like song bout length (Alward, Balthazart et al. 2013). Somewhat surprisingly, hypothyroidism did not affect the majority of neuroplasticity changes, only the myelination of several tracts during the photostimulated phase were affected. However, it is important to note that even low concentrations of testosterone appear sufficient to induce seasonal neuroplasticity in songbirds (Tramontin, Perfito et al. 2001, Caro, Lambrechts et al. 2005). The presence of a yellow beak in the MMI treated birds meant that there was still some testosterone in the circulation (Ball and Wingfield 1987), presumably enough to induce seasonal neuroplasticity. Furthermore, plasma T3 concentrations were negatively correlated to the fractional anisotropy changes in several main song control nuclei, which is in line with the finding that both control and hypothyroid starlings showed decreased thyroid hormone levels during the photosensitive period. We hypothesize that a global reduction of circulating thyroid hormones might be necessary to lift the brake imposed by the photorefractory period, whereas local fine-tuning of thyroid hormone and testosterone concentration through an upregulation of thyroid hormone transporters LAT1 and aromatase activity respectively could activate certain genes and mechanisms associated with neuroplasticity. However the effects of thyroid hormones are complicated by their effect on the HPG-axis and the difference between circulating and intracellular thyroid hormone levels. Therefore, this study is just one of the first steps to disentangle the influence of thyroid hormones on seasonal neuroplasticity and provides a framework for future studies to further investigate the molecular changes induced by thyroid hormones in a seasonal songbird.

### 4.1 Expression of thyroid hormone inactivating enzyme *DIO3* and thyroid hormone transporter *LAT1* in HVC change over photoperiods

We established the seasonal expression of several thyroid hormone regulating genes in the song control system using ISH, which is represented in a schematic overview in **Figure 10**. Especially the HVC experiences seasonal differences in thyroid hormone regulating gene expression, namely for *DIO3* and *LAT1*. We validated that our cell counting method was accurate (correlation between cell count and stained surface was > 0.9 for both *DIO3* and *LAT1*) and that these expression levels where not caused by changes in cell density (via Nissl staining **Figure 3 – figure supplement 1**). During the photosensitive phase circulating thyroid hormones are low, but a local upregulation of thyroid hormone transporter *LAT1*, and downregulation of *DIO3* at the level of the HVC, suggests that the local thyroid hormone concentration in the HVC is higher during the photosensitive phase. The downregulation of *DIO3* is not matched by an upregulation of *DIO2*, since there is an apparent lack of *DIO2* expression in the entire telencephalon. All vertebrates need at least a minimum of brain *DIO2* activity to survive but research in chicken and zebra finches already indicated that *DIO2* expression in the brain decreases with age (Van Herck, Delbaere et al. 2015, Raymaekers, Verbeure et al. 2017). Our data seem to confirm that at least in adult birds, *DIO2* expression is too low to be detected by our ISH staining method. This is in stark contrast with what was found during developmental neuroplasticity in the song control nuclei in zebra finches, where the expression of *DIO2* remained high during the sensitive and sensorimotor learning phases (Raymaekers, Verbeure et al. 2017). Several other studies have further suggested that local presence of DIO2 is essential for critical period learning or that correct local balancing of active THs is essential in plastic brain areas involved with learning (Guadaño-Ferraz, Escámez et al. 1999, Yamaguchi, Aoki et al. 2012). In zebra finches DIO2 was most intensively expressed in endothelial cells of blood vessels. Potentially, angiogenesis and hormone supply via the blood-brain barrier are more important and pronounced during developmental neuroplasticity than for adult seasonal neuroplasticity.

**Figure 10:**
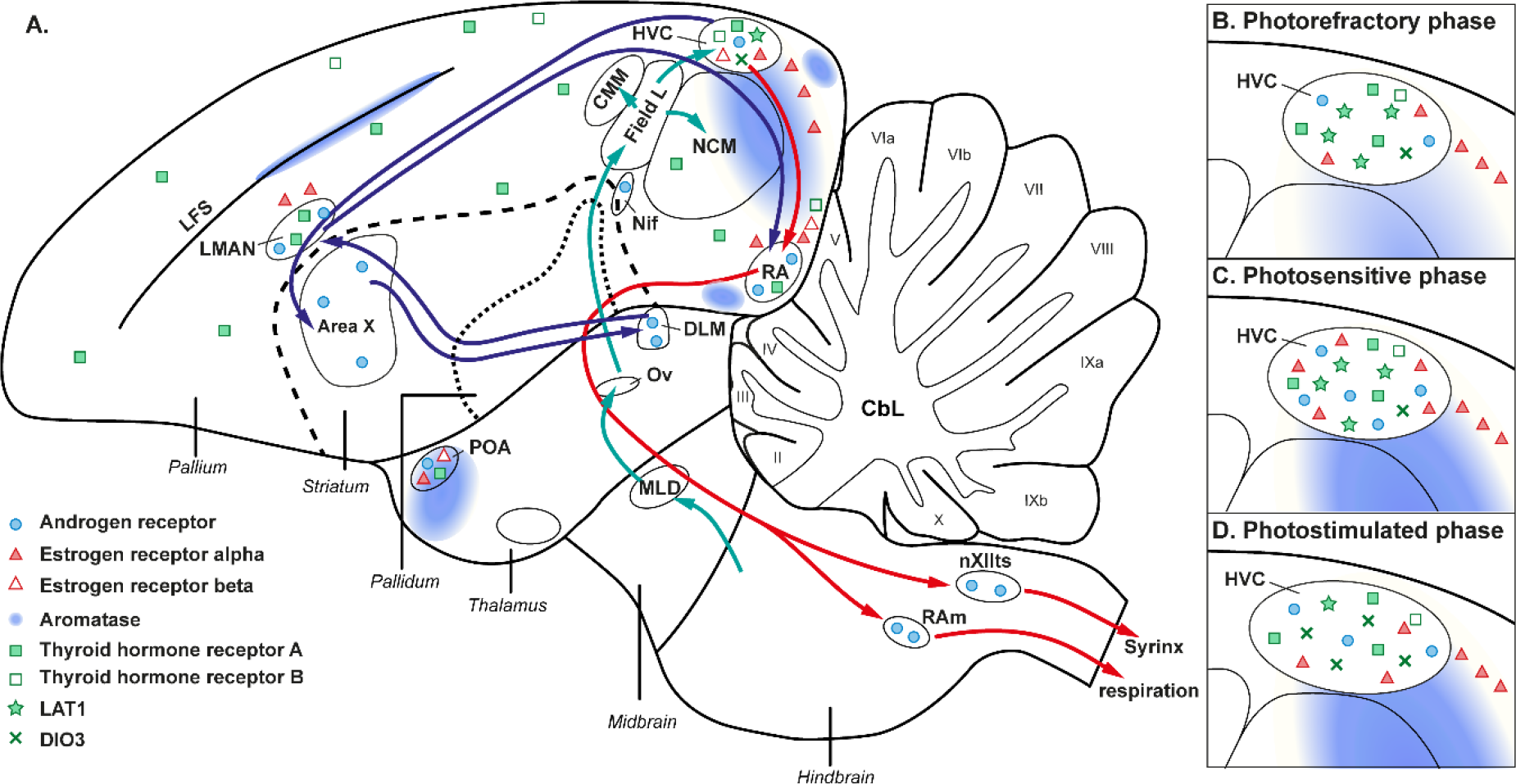
Schematic overview of the steroid and thyroid hormone receptors within the songbird brain (A) and how this expression changes seasonally at the level of the HVC (B - D). Androgen receptor, estrogen receptor and aromatase expression is based on prior studies in starlings and canaries (Gahr 1990, Bernard, Bentley et al. 1999, Metzdorf, Gahr et al. 1999). The expression of thyroid hormone regulating genes is derived from our own results. (B-D) In canaries the estrogen and androgen receptor expression is higher during photosensitive and early photostimulated phase compared to the photorefractory state (Gahr and Metzdorf 1997, Fusani, Van’t Hof et al. 2000). In starlings aromatase activity and androgen receptor density was the highest during the photosensitive phase (Riters, Baillien et al. 2001) (Riters, Eens et al. 2002).

Interestingly, several microarray studies in other seasonal songbirds have shown *DIO2* expression in HVC and RA with seasonal changes in expression. Stevenson, Replogle et al. (2012) found that in starlings the *DIO2* expression in RA was the highest during the photosensitive phase, which is in line with our hypothesis of local upregulation of thyroid hormone regulating genes in the song control system during the non-breeding periods. There are however some species differences as Gambel’s white crowned sparrows have upregulated *DIO2* expression during breeding conditions in HVC and RA relative to short day conditions (Thompson, Meitzen et al. 2012).

The inverse ratio between thyroid hormone transporter *LAT1* and *DIO3* is seen during the photostimulated phase. Even though circulating T4 levels are elevated during this phase, the influx of thyroid hormones and the potential activity of T3 seems to be repressed in the HVC. Local DIO2 and DIO3 activity can fine-tune intracellular thyroid hormone concentration at least partially independent of the circulating thyroid hormone levels. For example in Japanese quail, reciprocal switching of *DIO2* and *DIO3* expression upon photostimulation can locally increase T3 concentration by a 10-fold, even though plasma concentrations do not change (Yoshimura, Yasuo et al. 2003). Interestingly, this could mean that, whereas increased uptake of thyroid hormones by *LAT1* induces neuroplasticity, *DIO3* activity is a possible factor for stability of neural tissue as would be expected in the photostimulated stage.

Apart from DIO3 and LAT1, our study also looked at the expression of thyroid hormone receptors and other thyroid hormone transporters. Surprisingly, other thyroid hormone transporters like *MCT8, MCT10* or *OATP1C1,* were not detected by ISH in any photoperiod in the starling’s telencephalon, even though MCT8 is the transporter with the highest affinity for thyroid hormones (Bourgeois, Van Herck et al. 2016). Several studies, in mammals as well as birds, have shown that MCT8 is highly expressed in the developing brain and is considered as the most important thyroid hormone transporter during developmental neuroplasticity (Van Herck, Geysens et al. 2013, Raymaekers, Verbeure et al. 2017). However, our data suggest that the situation might be different in adult birds, where *LAT1* seems to be more prominent as a mediator in seasonal neuroplasticity.

We found *THRA* expression in large parts of the pallium including HVC, RA and LMAN, whereas the striatum including Area X lacked *THRA* expression, which is in line with the expression pattern of *THRA* in adult zebra finches (Raymaekers, Verbeure et al. 2017). In contrast, *THRB* was not detected in any song control nucleus. These observations align with what has been observed in chickens: *THRA* expression seems to be widespread while *THRB* is more locally expressed and exerts some specific functions for brain development (Forrest, Hallböök et al. 1991, Bradley, Towle et al. 1992, Darras, Van Herck et al. 2011, Van Herck, Geysens et al. 2013). Interestingly the expression pattern of *THRA* and *THRB* does not change seasonally, indicating that while at least *THRA* is necessary for proper genomic thyroid hormone action in the song control system, the level of thyroid hormone receptor expression is not a regulatory factor of song control system plasticity.

Together these findings suggest an active role for thyroid hormones during the non-breeding phase to induce plasticity in the song control system, in contrast to the photostimulated phase where thyroid hormones are locally decreased in the HVC to promote stabilization of the song **(Figure 12)**.

### 4.2 Thyroid hormones regulate HPG-axis activity in starlings

In the second experiment, we deprived the starlings of circulating thyroid hormones by MMI treatment during the photosensitive and photostimulated phase to investigate their effect on testosterone, song and neuroplasticity. MMI is an anti-thyroid drug inhibiting the synthesis of T3 and T4 at the level of the thyroid gland, thereby gradually depleting circulating thyroid hormone levels. MMI treatment was started on the first short day of the photosensitive period, as earlier treatment could terminate the photorefractory period (Dawson, Goldsmith et al. 1985). After 4 weeks of MMI treatment thyroid hormone levels were successfully depleted, which is consistent with the findings in Japanese quail, where MMI treatment for 4 weeks resulted in decreased thyroid hormone levels, testes weight and testosterone levels (Weng, Saita et al. 2007). In immature rats MMI affected the HPG-axis, decreasing luteinizing hormone (LH) and testosterone plasma concentration, but also decreased the testes binding sites for LH by 30% (Valle, Oliveira-Filho et al. 1985, Pérez, Meddle et al.

2018). The present results show that hypothyroidism also inhibited the photostimulation-induced increase in testosterone in starlings. Yoshimura, Yasuo et al. (2003) demonstrated that in Japanese quail thyroid hormones play a crucial role in the activation of the HPG-axis upon photostimulation. Exposure to long days increases hypothalamic T3 concentration as a result of a local upregulation of *DIO2* and downregulation of *DIO3* in the mediobasal hypothalamus. This locally produced bioactive T3 subsequently results in morphological changes in GnRH nerve terminals enabling GnRH secretion resulting in activation of the HPG-axis (Yoshimura 2013). In starlings under natural photoconditions, this phenomenon was observed to a lesser extent, but it did not corroborate the proposed photoperiodic regulation of the HPG-axis through *DIO2/DIO3* expression (Bentley, Tucker et al. 2013). It is suggested that both phylogenetic differences and the acuteness at which photoperiods are changed in experiments might influence the changes in hypothalamic thyroid hormone regulation. Likely, the regulatory mechanisms in natural conditions are similar but more gradual. In agreement with manipulation studies in quail where the DIO2 inhibitor iopanoic acid prevents testicular growth (Yoshimura, Yasuo et al. 2003), our manipulation study confirms the active role of thyroid hormones in the activation of the HPG-axis **(Figure 12)**.

### 4.3 Hypothyroidism affects song behavior by depleting testosterone

Of particular interest is the relationship between hormonal changes, song behavior and neuroplasticity as they change over the different seasonal states that were induced by the photoperiodic manipulations (schematic overview in **Figure 11**). By using *in vivo* techniques such as MRI, we established that song bout length is positively correlated to the testosterone levels, beak color and neuroplasticity associated fractional anisotropy changes (**Figure 11**). The correlation between song bout length and testosterone is only present in the control group, since MMI treatment inhibited the photostimulation-induced increase in testosterone. However, beak color was still correlated to the song bout length in the MMI group, indicating that even low concentrations of testosterone, below detection limit of the RIA assay but enough to induce beak color changes, are linked to changes in song bout length. The neural substrate of song bout length includes, as expected, song control nuclei like HVC, RA and LMAN, which concurs with earlier correlation studies in starlings and other songbirds (Bernard, Eens et al. 1996, Garamszegi and Eens 2004). However, also other structures like the lateral septum and cerebellar lobule VI-VII are correlated to song bout length. At the level of the lateral septum Merullo, Cordes et al. (2015) showed a positive correlation between neurotensin (a neuropeptide that regulates dopamine activity) expression and both sexually motivated song and non-vocal courtship behaviors in starlings. Whereas in songbirds, only few studies looked at the role of the cerebellum in song behavior through the cerebello-thalamic-basal ganglia pathway (Person, Gale et al. 2008, Nicholson, Roberts et al. 2018, Pidoux, Le Blanc et al. 2018), in humans, the cerebellum is known to be involved in language tasks (lobule VI and crus I/II) and phonological storage during the maintenance of verbal information (VIIb/VIII). Prior lesion study showed that lesioning Area X caused structural remodeling of the cerebello-thalamic-basal ganglia pathway extending into cerebellar lobules VIb and VII (Hamaide, Lukacova et al. 2020). Our results further corroborate that cerebellar lobules VI-VII retain the neural substrate for song behavior. Furthermore, these parts of the cerebellum show seasonal neuroplasticity, both in male and female starlings (Orije, Cardon et al. 2021).

**Figure 11:**
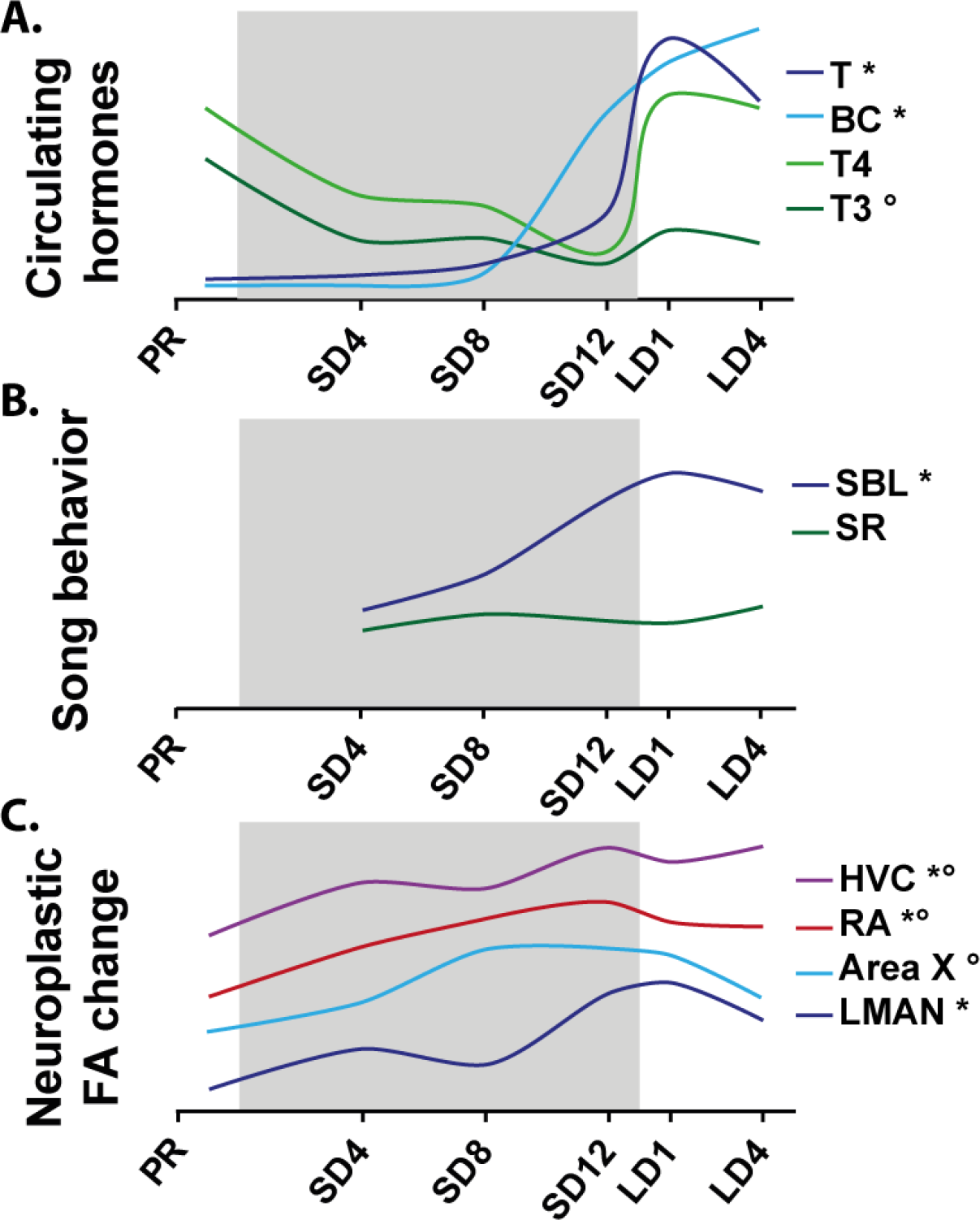
Schematic overview of the seasonal changes in circulating hormones, thyroid hormone (TH) and testosterone (T), song behavior and neuroplasticity changes in fractional anisotropy (FA). Correlations to song bout length (SBL) and T3 are indicate by * and ° respectively. The grey area indicates the photosensitive period. Abbreviations: BC, beak color.

The absence of high testosterone levels in hypothyroid starlings during the photostimulated phase could explain some of the differences in song behavior. Even though song rate did not change significantly over time in both groups, hypothyroid starlings sang with a lower song rate during photostimulation compared to the control group. Testosterone is known to increase the motivation to sing, by interacting with the medial pre-optic nucleus (POM), since local testosterone implantation in the POM of castrated male canaries increased their song rate (Alward, Balthazart et al. 2013).

Interestingly, hypothyroid starlings keep on increasing their song bout length during the photostimulated phase, compared to the control group, which sing at a stable song bout length during photostimulation. This lack of stability in the song bout length during photostimulation could be due to the absence of high testosterone concentrations, as testosterone is known to be necessary for song crystallization and stabilization in sparrows, zebra finches and canaries (Marler, Peters et al. 1988, Korsia and Bottjer 1991, Williams, Connor et al. 2003). Blocking testosterone by castration or by the androgen receptor antagonist flutamide, causes variability in song of zebra finches and canaries (Wang, Liao et al. 2014, Alward, Balthazart et al. 2017). More specifically blocking androgen receptors in the RA of male canaries, caused song duration to increase, whereas specific blocking in HVC increased the variability in used syllables (Alward, Balthazart et al. 2017) **(Figure 12)**.

**Figure 12:**
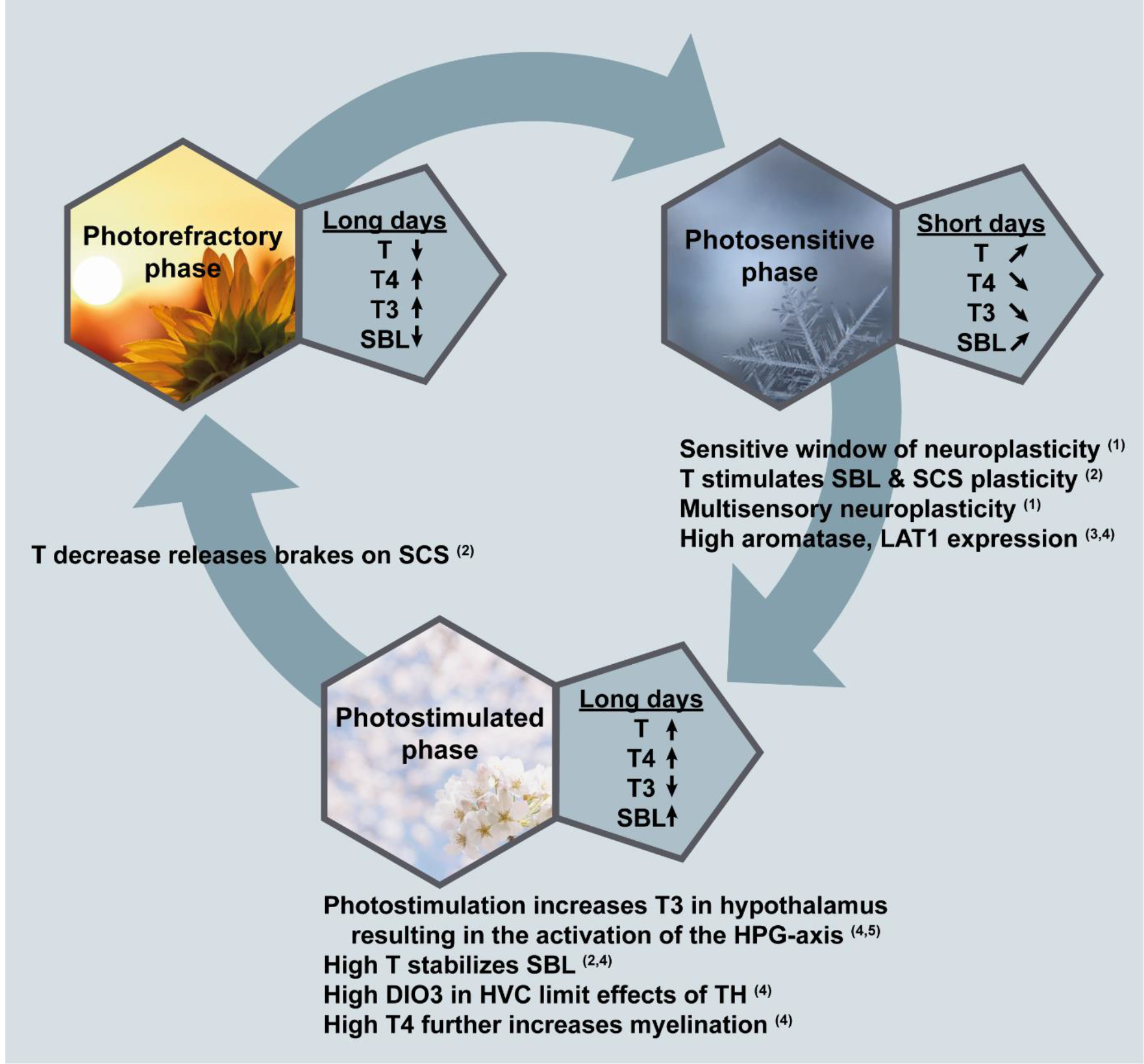
Schematic overview summarizing the seasonal changes in hormones, song behavior and neuroplasticity found in the current study and the hypotheses derived from this and prior studies. For each seasonal stage the plasma hormone concentrations (testosterone (T), thyroid hormones (TH) thyroxine (T4) and triiodothyronine (T3)) and song bout length (SBL) as a read out for song quality were summarized. During the short days of the photosensitive period, plasma T concentrations and SBL gradually increase, while plasma T4 and T3 concentrations gradually decrease. This period was demonstrated to be a sensitive window for increased neuroplasticity in the song control system (SCS), all sensory systems and the cerebellum (see (1) (Orije, Cardon et al. 2021)). During this period, T can induce increased SBL and SCS neuroplasticity (see (2) Orije, Cardon et al. (2020)). Even though the circulating concentrations of T and thyroid hormones are low, their effects might be maximized by local fine-tuning through upregulation of aromatase in the telencephalon (see (3) Riters, Baillien et al. (2001)) and thyroid hormone transporter LAT1 in HVC (see **Figure 2** of this study (4)). Upon long days during photostimulation, T3 levels increase in the hypothalamus resulting in the activation of the HPG-axis (see (5) Yoshimura (2013)). The resulting peak in T levels coincided with high SBL and T4 levels. These high T levels help to stabilize the SBL during this phase. High DIO3 expression in the HVC represses the potential activity of T3 or T4, preventing further neuroplasticity in order to promote song stability (see **Figure 2** of this study (4)). In other parts of the brain, high T4 is required for myelination and hence consolidation of several white matter tracts that were modulated and shaped in the photosensitive stage (see **Figure 7** of this study (4)). Continuation of long days eventually leads to the photorefractory phase, where T levels decrease again due to gonadal regression, thereby releasing again the brakes of neuroplasticity on the SCS and allowing the song to become plastic again (see (2) Orije, Cardon et al. (2020)).

Many early studies concluded that relatively high testosterone concentrations seem to be involved in high song rate and song stability during the photostimulated phase or the annual cycle. More recent studies have found that low concentrations of testosterone are sufficient to induce seasonal neuroplasticity (Tramontin, Perfito et al. 2001, Caro, Lambrechts et al. 2005). In our study, changes in beak color represented low concentrations of testosterone that are biologically active and correlated to song bout length, which has its neural substrate in several song control nuclei, lateral septum and cerebellum.

### 4.4 Complex interaction between thyroid hormones and neuroplasticity

Deprivation of thyroid hormones during the photosensitive period did not prevent neuroplasticity from occurring. In contrast, certain regions within the MMI group showed earlier significant differences in FA during the short days compared to the control group, pointing to an earlier onset of neuroplasticity coinciding with the earlier depletion of T3 and T4. Under natural conditions, the level of circulating thyroid levels is low during the photosensitive period. By administrating MMI, we rapidly decrease circulating thyroid hormone levels to a minimum. The decrease of circulating thyroid hormones is necessary to terminate the photorefractory period for the reproductive system (Dawson, Goldsmith et al. 1986). The photorefractory period depends on thyroid hormones to prevent reproductive maturation in response to long day length (Nicholls, Goldsmith et al. 1988).

Photorefractoriness not only forms a brake on gonadal maturation but also prevents an increase in singing activity in response to testosterone treatment, in contrast to photosensitive female starlings treated with testosterone (Rouse, Stevenson et al. 2015). Potentially, the global reduction of circulating thyroid hormones is necessary to lift the brake on singing and on reproduction imposed by the photorefractory period. Interestingly, the circulating T3 level presents a negative within-subject correlation to neuroplasticity in HVC, Area X and RA surroundings both in the control and hypothyroid group, indicating that decreasing circulating thyroid hormone levels are associated with the neuroplasticity during the photosensitive phase. This negative correlation between T3 and neuroplasticity seems in contrast to the thyroid hormone regulating genes in HVC, which supports an active role of thyroid hormones during the photosensitive phase. We hypothesize that while a global reduction in thyroid hormones is necessary to lift the brake imposed by the photorefractory period, local fine-tuning of thyroid hormone concentration through thyroid hormone regulating proteins like LAT1 and DIO3 could activate mechanisms associated with neuroplasticity. A similar local fine-tuning of testosterone levels during the photosensitive phase has been suggested by Riters, Baillien et al. (2001). They found increased aromatase activity and decreased 5β-reductase activity during the photosensitive period in male starlings, increasing the brain’s sensitivity to testosterone despite the low circulating testosterone levels (**Figure 10**).

Prior studies have shown that the photosensitive phase acts as a sensitive window for multisensory neuroplasticity (Cornez, Collignon et al. 2020, Orije, Cardon et al. 2021). Multiple studies looked at potential causes for the reopening of a critical or sensitive period and there seem to be two main components involved: (1) the balance between the excitatory-inhibitory (E-I) circuits and (2) the molecular brakes that are imposed to limit adult neuroplasticity. After an initial critical period, neuroplasticity can be reintroduced when the E-I balance shifts more towards the excitation rather than inhibition due to increased excitation or reduced inhibition. Whether and how thyroid hormones directly partake in the regulation of molecular mechanisms of reopening a sensitive window in vocal learning remains to be elucidated. Thyroid hormone function is complex, some genes are upregulated in the presence of T3 (positively regulated genes), whereas some genes are down-regulated (negatively regulated genes). Gil-Ibanez, Garcia-Garcia et al. (2017) showed that both positively and negatively regulated genes can play a role in neuroplasticity. Further studies are required to identify the genetic expression changes upon thyroid hormone manipulation in starling brain, and to establish how these genetic changes affect neuroplasticity.

At the end of the photosensitive phase, the sensitive window closes by imposing molecular brakes like PNN and myelin. PNNs are highly expressed in the HVC of adult zebra finches after their song learning and predicts song maturity (Balmer, Carels et al. 2009). However, starlings have far less PNNs, compared to closed-ended learning zebra finches, suggesting that PNN might play less of a role in the control of seasonal neuroplasticity of open-ended learning starlings (Cornez, Madison et al. 2017). Our prior studies suggested myelination rather than PNNs as another candidate mechanism that contributes to ending a sensitive window in open-ended learners like starlings (Orije, Cardon et al. 2021). In the control group, several tracts including OM, TrO and tFA increased in fractional anisotropy upon photostimulation, due to decreased radial diffusivity, which implies an increase in myelination. This increased fractional anisotropy was absent in the MMI-treated group, which could reflect the thyroid hormone effect on myelination. Myelin basic protein is a positively regulated thyroid hormone dependent gene, which could respond to the normally increased T4 plasma concentrations during the photostimulated phase. Furthermore, thyroid hormone therapy was shown to induce myelinogenesis in an animal model of demyelination (Harsan, Steibel et al. 2008). So increased levels of testosterone and thyroid hormone levels during photostimulation could play a role in ending a sensitive window for plasticity in song behavior and its underlying neural structure through myelination acting as a molecular brake **(Figure 12)**.

### 4.5 Future perspectives

Methimazole treatment did not only drastically reduce thyroid hormones to a minimum, it also inhibited the increase of testosterone upon photostimulation. In the current study we altered both thyroid and testosterone concentrations. Future studies might want to control the amount of circulating testosterone by castration or testosterone implantation during photostimulation, to be able to solely investigate the effects of thyroid hormone manipulation. This way the suggested effect of thyroid hormone on myelination of several tracts during the photostimulated phase could be attributed to the lack of thyroid hormone’s positive effect on myelination, if myelination is still affected with MMI treatment combined with testosterone supplementation during photostimulation.

Importantly, methimazole treatment does not completely abolish thyroidal hormone production and thus low concentrations of thyroid hormones could still be locally picked up within the brain, as suggested by the increased LAT1 expression in the HVC. Future studies could consider other manipulations to investigate the effects of thyroid hormones on neuroplasticity in songbirds and their song behavior. For example, iopanoic acid both inhibits thyroid hormone release from the thyroid gland but also the conversion of T4 to T3 by inhibiting DIO2, but it is less practical for long term treatment strategies (Pant and Chandola-Saklani 1993). In the past both surgical and radioactive thyroidectomy have been applied to study the effects of thyroid hormones on seasonal reproduction (Dawson, Goldsmith et al. 1986, Bentley, Dawson et al. 2000). Analogous to experiments examining the local effects of testosterone by local androgen receptor inhibition (Alward, Balthazart et al. 2017), thyroid hormone antagonists and thyroid hormone receptor modulators have been developed and could be injected locally within specific song nuclei, to investigate their role in song behavior (Lim, Nguyen et al. 2002, Yoshihara and Scanlan 2003, Raparti, Jain et al. 2013).

## 5 Material & Methods

### 5.1 Subjects & Experimental setup

Wild European starlings (Sturnus vulgaris) used in the *in situ* hybridization experiment were caught in Baltimore, USA during mid-late winter 2013 and subsequently treated by the research group of prof. Gregory Ball. 24 male adult starlings were divided in 3 groups of 8, and treated in order to represent the photosensitive, photostimulated and photorefractory states. All birds were initially maintained on short day lengths (8 hours of light, 16 hours of dark, 8L:16D) to keep them in a photosensitive state. Photoperiod manipulations were applied over a period of 12 weeks. Group 1 was immediately put on long day lengths (14L:10D) for 12 weeks so they would be photorefractory by the end of photoperiod manipulation. Group 2 was kept on 8L:16D for 12 weeks so that they would remain photosensitive. Group 3 was kept on 8L:16D for 8 weeks, then switched to long day lengths (14L:10D) for 4 weeks to make them photostimulated. In order to control for handling stress associated with transferring group 2 to a different photoperiod, all groups were transferred into a new cage located within the same room. Birds remained with the same cage mates throughout the entire experiment. No females were present in any group. While studies showed that the presence of females influences neurological attributes and singing rate in canaries (Alward, Mayes et al. 2014, Shevchouk, Ball et al. 2017), and even slightly increases the growth of the sparrow HVC and RA compared to isolated birds, photoperiod is still the main instigator of song control nucleus growth (Tramontin, Wingfield et al. 1999).

Food and water was supplied *ad libitum*, and supplementary spinach was provided daily. In order to verify sex and measure testis growth, exploratory laparotomies were performed prior to placing birds into their experimental groups. Under isoflurane anesthesia, a small incision was made on the bird’s left side, and calipers were used to measure the testis diameter. The incision was closed with nylon sutures, and birds were allowed to recover for one week. After one week, they were placed into the appropriate experimental groups. At the end of the 12 week period, all birds were anesthetized via injection of secobarbital followed by cardiac perfusion with 4% paraformaldehyde. Brains were dissected out and placed into 4% paraformaldehyde. The testes were dissected out, measured with calipers and weighed. Bird body weights were recorded, and the color of their beaks was scored to monitor physiological response to the photoperiod. All procedures were in accordance with the animal welfare regulations of the John Hopkins University of Baltimore, Maryland, USA. Brains were quickly frozen, stored at -80 °C and sent to the research group of Comparative Endocrinology at the KU Leuven together with blood samples, where all subsequent procedures took place.

The second experiment was performed on thirty male starlings (*Sturnus vulgaris*) that were wild caught as adults in Normandy (France) in November 2014. All animals were housed in two large indoor aviaries (L x W x H: 2.2 x 1.4 x 2.1 m) at the University of Antwerp with food and water ad libitum with artificial light dark cycle. Starting from January 2013, all birds were kept in a long day photoperiod (16L/8D) in order to remain photorefractory. The housing and experimental procedures were performed in agreement with the Belgian and Flemish laws and were approved by the Committee on Animal Care and Use of the University of Antwerp, Belgium (2014-52).

Starlings were divided into two groups: a hypothyroid group (N=16) and a control group (N=14). The study was started when all birds were photorefractory. Then they were switched from long to short (8L:16D) days to induce the return to photosensitivity. Methimazole (MMI) treatment was started in one group to induce hypothyroidism. By supplementing the drinking water with 0.05% MMI, the endogenous stock of THs gradually decreased until it was fully depleted after 2-3 weeks. MMI treatment was continued for the remainder of the experiment. After 12 weeks of short days, the photosensitive starlings were switched back to long days (16L:8D) so that they became photostimulated. In parallel the control group was exposed to the same photoperiodic regime without receiving any hormone manipulation.

We monitored the neuroplasticity repeatedly at 6 different time points. The first time point was at the end of the photorefractory state (PR). After switching to short days we measured every 4 weeks to follow up the song control system plasticity during the photosensitive period (SD4, SD8, SD12). Additionally, we measured after 1 week of long days, as it is known that exposure to 1 long day can already affect TH.

Finally, we measured after 4 weeks on long days (LD4), when starlings were fully photostimulated. At each time point, songs were recorded, blood samples were taken, and MRI (DTI and 3D) was acquired. In addition body weight and beak color were registered.

### 5.2 *In situ* hybridization

30 μm thick coronal brain sections were made using a cryostat and kept free floating in cryobuffer (1.59 g/l NaH2PO4, 5.47 g/l Na2HPO4, 300 g/l sucrose, 10 g/l polyvinylpyrrolidone and 300 ml/l ethylene glycol in RNase free H2O) in 24 series so that each subsequent section in a series was 720 μm apart from the previous one. Sections were stored in cryobuffer at -20°C until used.

#### 5.2.1 DNA cloning and probe synthesis

DNA cloning and probe synthesis of a partial sequence of the target genes in starling was done similarly to the method used for zebra finch genes (Raymaekers, Verbeure et al. 2017). RNA was isolated from the brain of an adult male starling. The primers used for cloning are shown in **Table 4**. Cloning was successful except for the sequence of OATP1C1. Therefore, we instead used the corresponding probe for zebra finch (Raymaekers, Verbeure et al. 2017) that has 94% sequence homology with the corresponding starling sequence, which is sufficient to generate specific signal.

**Table 4:**
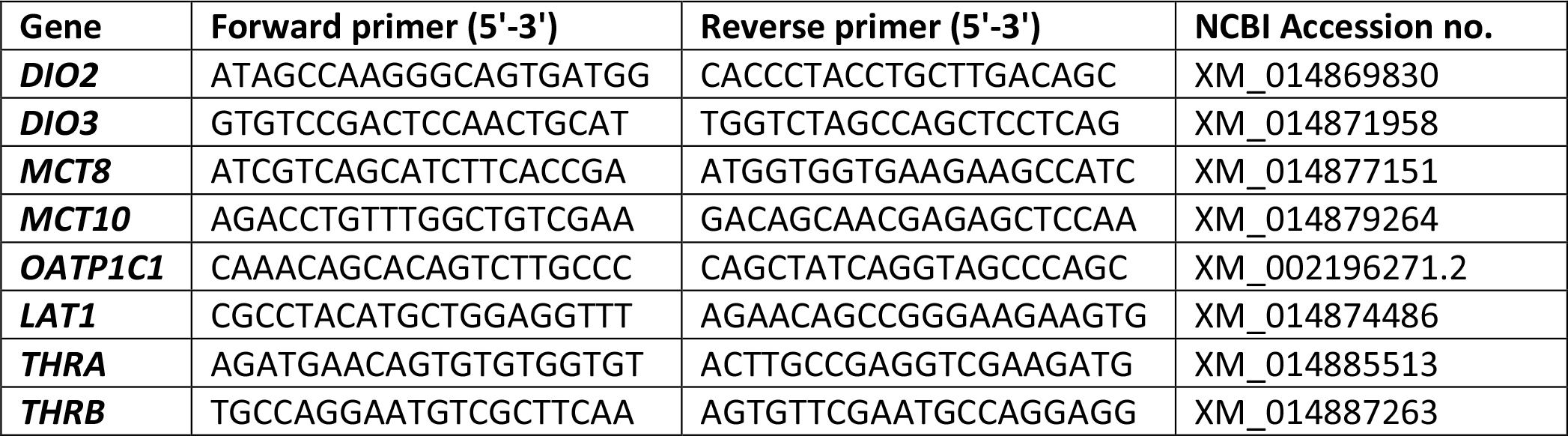
Primers for cloning of partial sequences of thyroid hormone regulators in starling.

#### 5.2.2 *In situ* hybridization staining

The ISH protocol used for detection of thyroid hormone regulator genes in starling closely resembled the protocol used for zebra finch (Raymaekers, Verbeure et al. 2017). From each series, sections containing song control nuclei were selected and placed individually in wells of a 24-well plate. This was on average 3 sections for Area X, 2 for LMAN, 2 for HVC and 2 for RA. Of each photoperiod, one brain was omitted from staining due to cutting or freezing damage; thus the number of birds (N) stained for each photoperiod was 7. ISH was performed with both anti-sense and sense (negative control) probes.

#### 5.2.3 Nissl staining

Of 3 birds of each photoperiod, 2 HVC sections, 720 µm apart, were stained with cresyl violet. Sections were rinsed in PBS, incubated in 1% cresyl violet in H20 for 1 min and then rinsed three times in H20 containing 2.5% acetic acid for 1 min. Sections were again rinsed in H20, mounted onto glass slides, incubated in xylene for 30 seconds and coverslipped with DPX (Sigma).

#### 5.2.4 *In situ* hybridization imaging and data analysis

Photomicrographs in the regions of interest were taken for all markers using a stereomicroscope (Carl Zeiss). Data in HVC from the *DIO3* and *LAT1* ISH and the Nissl staining were quantified (on average 2 sections per bird for each marker) using Fiji software (ImageJ, National Institute of Health) and statistically analyzed using GraphPad Prism 5 (GraphPad Software, Inc.). We only analyzed 1 side of each section, selecting a 100 000 μm² square box was selected in the middle of HVC (central on the mediolateral and dorsoventral axes through the nucleus). Inside that box, *DIO3+* cells, *LAT1+* cells and cells stained by cresyl violet were counted. Cells were counted as a ’stained cell’ if the dark signal was delineated in the form of at least a semicircle (180 degrees or more) around the center. Cell counts of both sections from one bird were then averaged to obtain 1 numerical value per bird. Signal throughout HVC was generally homogeneously spread, but for rare sections with heterogeneous signal, larger areas were analyzed and data normalized. We analyzed the differences in *DIO3+, LAT1+* or Nissl stained cells between photoperiods using a linear mixed model. When appropriate, further comparisons were performed with the Bonferroni *post hoc* test or Dunn’s *post hoc* test respectively to analyze differences between two photoperiods.

To further evaluate the validity of the cell count method used for the *DIO3* and *LAT1* ISH, 10 random sections of both ISHs were analyzed using the semi-quantitative manner of ’stained surface fraction’ as used for the zebra finches (Raymaekers, Verbeure et al. 2017). For each section, a 100 000 μm² square box was selected in the middle of HVC (central on the mediolateral and dorsoventral axes through the nucleus). The image was converted to greyscale (each pixel is assigned a brightness value between 0 and 255) and a brightness threshold was manually set for each image so that dark staining was counted as ’stained’ while background was not. This way, the stained surface area was calculated and divided by the total area of the 100 000 μm² box. The stained surface fraction values were plotted against their corresponding original cell count value. Linear regression analysis was performed and the coefficient of determination (r²) was calculated using Prism 5 to check for correlation.

### 5.3 Blood sample analysis

At each time point of the hypothyroidism experiment blood samples were taken to assay plasma levels of T3, T4 and testosterone. In order to limit variation of plasma levels due to the time of the day, blood samples were taken at each stage within a 2 hour window at noon between 11u30 and 13u30. Animals were already subdivided in small cages for individual song recordings. This allowed fast capture and blood sampling (within 5 min.) to minimize increase in testosterone as a result of acute stress (Van Hout, Eens et al. 2010). The alar wing vein was punctured with a 25-gauge needle to collect 300-500 µl of blood into heparin-coated capillary tubes. Blood samples were centrifuged for 10 min at 2060g while cooling to 4°C. After centrifugation, the plasma was collected and frozen at -20°C, where the samples remained until analysis. Radioimmunoassay for testosterone, T3 and T4 were performed as described previously in (Orije, Cardon et al. 2021) and (Reyns, Janssens et al. 2002) respectively.

### 5.4 Song analysis

Song was recorded with Behringer C-2 condensator microphones, connected to a PR8E multi-channel preamplifier (SM Pro audio) and ‘Delta Series’ Breakout box (M-audio). The microphones were placed on top of individual cages. Due to technical issues, songs of birds were lost at PR and SD12 in both groups and at SD4 in the MMI treated group. Since fully isolated starlings do not sing as they would do in group context, the individual cages were placed next to the aviary. Using SoundExplorer (Developed by RF Jansen, University of Amsterdam) sonograms were created of the recordings. Based on the intensity and the spectral definition of the song bouts, individual song could be differentiated from background noise of the aviary. In line with prior song processing (Eens 1997, Van Hout, Pinxten et al. 2012) we defined a song bout as a period of at least 5s of song with pauses no longer than 1.5s. Starling song consists out of 4 distinct phrases, which are mostly performed in a fixed order. The song often starts with one or several whistles (whistle phrase), followed by a section of complex phrases (variable phrases) and rapid series of clicks or rattles (rattle or click phrase), before ending the song with high frequency song elements (high frequency phrases). Song bouts containing at least three different phrases were labelled as ‘complete song bouts’. Analysis of the song bout length was only performed on complete songs. If there were no complete song bouts at a certain time point, the average song bout length of incomplete songs was calculated and taken as a measure of the evolution in song production.

At each photoperiod songs were recorded for 48 hours. The song bout length measured in seconds and singing rate (average minutes spent singing per hour) were quantified as a measure of the singing behavior. The song rate was calculated as the average number of complete song bouts per hour. Statistical analysis of testosterone levels and song behavior was performed using linear mixed model via JMP Pro 13. *Post hoc* tests were Tukey corrected for multiple comparison. Differences were considered significant for p<0.05.

### 5.5 MRI data acquisition

The birds were initially anesthetized using 2% Isoflurane (Isoflo ®, Abbot Laboratories Ltd.) in a mixture of 30% O2 and 70% N2 at a flow rate of 600 ml/min. Throughout the entire imaging procedure, respiration rate was monitored with a small pneumatic sensor (SA Instruments, NY, USA) positioned under the bird. Depending on the breathing rate, the anesthetic dose was lowered, ranging between 1% - 2% isoflurane. Body temperature was monitored with a cloacal temperature probe and kept within narrow physiological ranges (41.0 ± 0.2 °C) using a warm air system with a feedback unit (SA Instruments, NY, USA).

All MRI measurements were performed on a 7T horizontal MR system (Pharmascan 70/16 US, Bruker Biospin, Germany). Each imaging session started with a T2-weighted 3D anatomical RARE scan (TR: 2000 ms; TE: 11 ms; RARE factor: 8; zero-filled to a matrix of (256x92x64) with voxel resolution (0.089x0.25x0.25) mm³). Subsequently, a 4 shot SE-EPI DTI scan (TR: 7000 ms; TE: 23 ms; d 4ms, D 12ms; b-value 670 s/mm²; 60 diffusion gradient directions; spatial resolution: (0.179x0.179x0.35) mm³; 28 coronal slices) was acquired. After the imaging procedure, birds were left to recover in a warmed recovery box before returning to the aviary.

### 5.6 MRI data processing

Diffusion data was analyzed with the Diffusion II toolbox in SPM12 software (Statistical Parametric Mapping, http://www.fil.ion.ucl.ac.uk/spm/software/spm12/). In a first step, the diffusion weighted (DW) images were realigned to correct for motion, using a rigid registration between the b0 images. This motion correction was then extended to all DW-images. Next, the realigned DW-images were co-registered to the anatomical 3D image of each subject using normalized mutual information as the similarity metric. Earlier a 3D template was created from all the subject anatomical 3D scans using ANTs (Advanced Normalization Tools)(Avants, Yushkevich et al. 2010), providing a standardized neuroanatomical space of the female starling brain. This facilitated normalization of the individual 3D images to the template. The normalization parameters from the 3D images were applied to the co-registered DW-images. Subsequently, the diffusion information was reoriented to correct for changes due to realignment, co-registration and normalization. Diffusion tensor was estimated from the DW- data and DTI parameter maps were derived (i.e. fractional anisotropy (FA), mean diffusivity (MD), eigenvalue 1 (λ1) or axonal diffusivity and radial diffusivity (RD, mean of λ2 & λ3) maps). Finally, all DTI parameter maps were smoothed using a Gaussian kernel of double the voxel size (FWHM= 0.36 x 0.36 x 0.48 mm³). After every processing step, images were checked for errors due to excessive motion, failed normalization or distortions. Outliers were removed from further processing.

### 5.7 Statistical analysis DTI

The normalized DTI parameter maps were used for voxel based analysis and ROI based analysis. The voxel based analysis included one way ANOVA to look for significant changes over time and multiple regression to investigate correlations to song behavior and testosterone. We performed these analyses with a whole brain mask and subsequently with a mask for song control nuclei, auditory system and their tracts. This way we ensured not to miss significant changes in relevant regions that might be obscured by the stringent correction for multiple comparisons in whole brain analysis.

ROI’s defined on the starling atlas (De Groof, George et al. 2016) were transformed to match the template space. Using the MarsBaR toolbox of SPM, mean DTI values could be extracted from the different ROI’s in automated way for a normalized ROI based analysis (Brett, Anton et al. 2002).

In addition, individual DTI parameter maps were reconstructed without normalization to the template, so that changes in volume were not obscured by normalization. Using the neuroanatomical contrast of the fractional anisotropy maps we could delineate different ROI’s (telencephalon, RA and HVC-RA tract) using AMIRA software (De Groof, Verhoye et al. 2006). This way we could determine volume and DTI changes within these ROI’s. Linear mixed model with *post hoc* Tukey correction was performed on the ROI data using JMP Pro 13. Differences were considered significant for p<0.05.

## Acknowledgements

The MRI equipment was funded by the Hercules foundation (Belgium, grant agreement AUHA/012) under the promoter-ship of Annemie Van der Linden. This research was funded by a grant from the Research Foundation – Flanders (FWO, project Nr G030213N)) awarded to Annemie Van der Linden and Veerle Darras, and the Interuniversity Attraction Poles (IAP) (“PLASTOSCINE”: P7/17) to Annemie Van der Linden. Jasmien Orije is a PhD fellow of the FWO (Nr 1115217N) and Elisabeth Jonckers received Postdoctoral fellowships of the FWO (Nr 12R1917N).

## Supplementary information

**Figure 3 – figure supplement 1:**
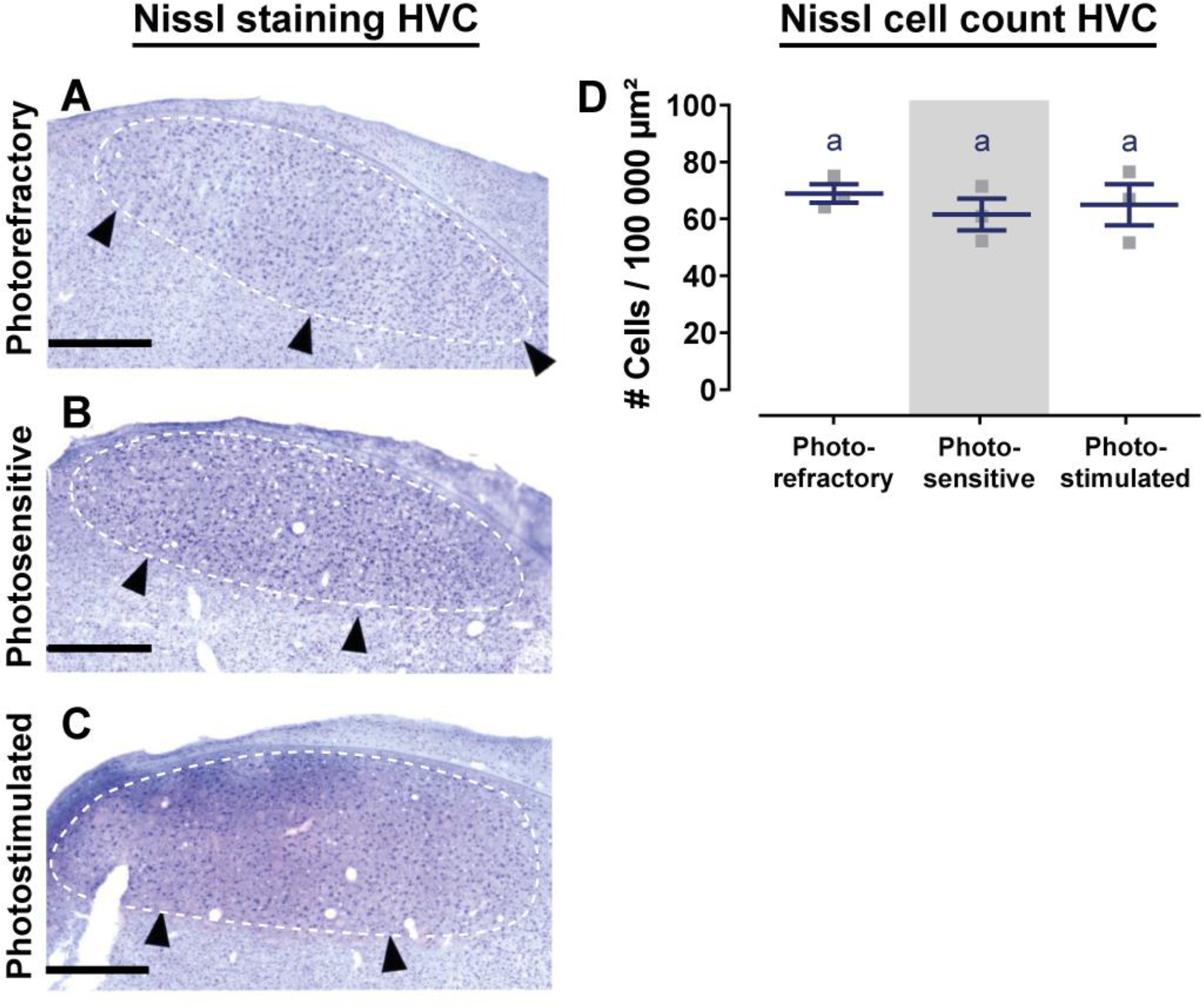
Representative images of the HVC of starlings in different photoperiodic states stained by cresyl violet (A, B, C) and quantitative analysis of the Nissl cell count in the HVC. The border of HVC is designated by white dashed lines and black arrowheads. Scale bar = 500 μm. Horizontal bars represent the average with standard deviation error bars (n=3). The grey area indicates the photosensitive phase.

**Figure 6 – figure supplement 1:**
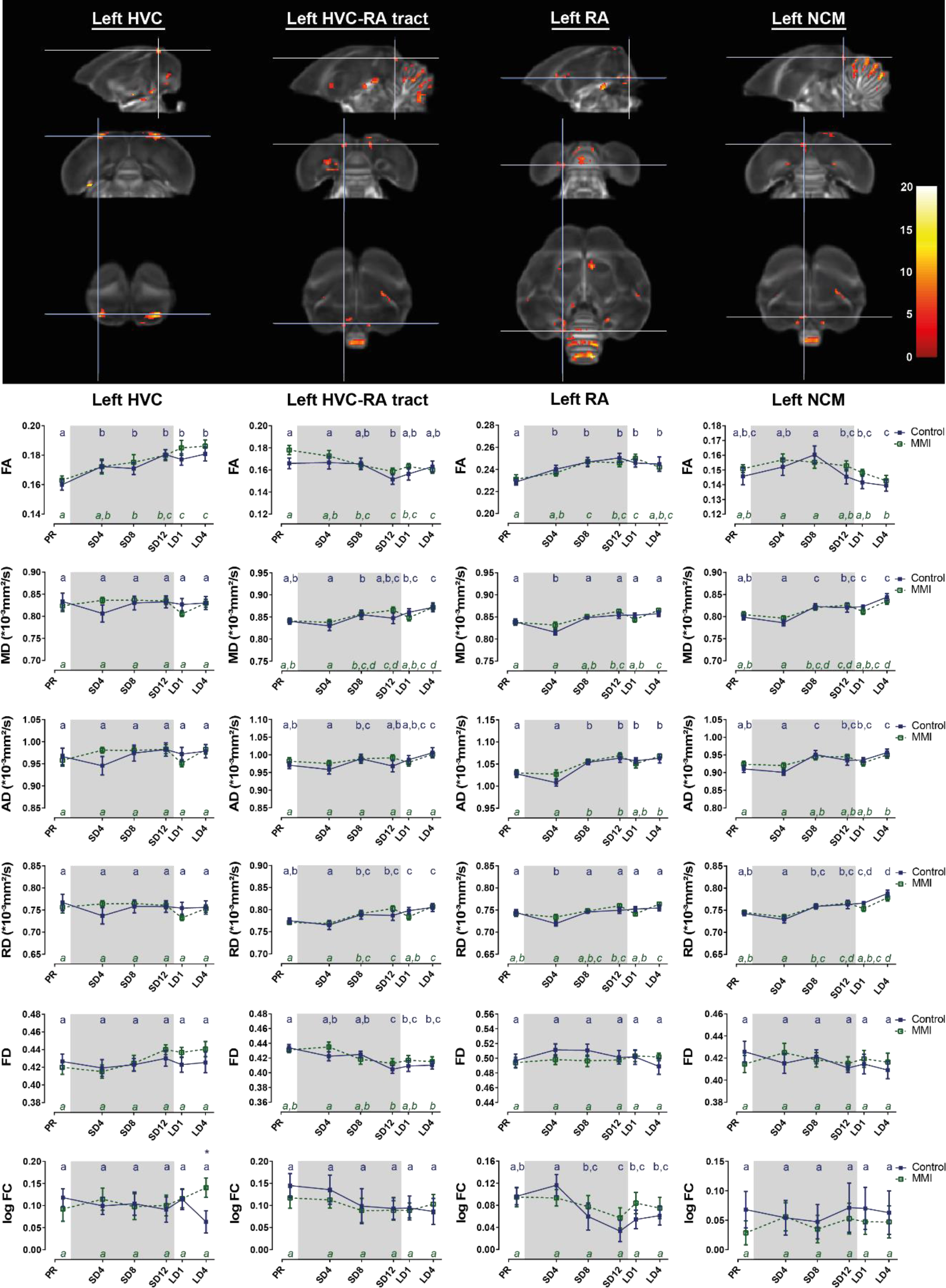
Summary of the significant longitudinal changes over time in FA, MD, AD, RD, FD and log FC extracted from ROI-based clusters at level of the left HVC, HVC-RA tract, RA and NCM.

**Figure 6 – figure supplement 2:**
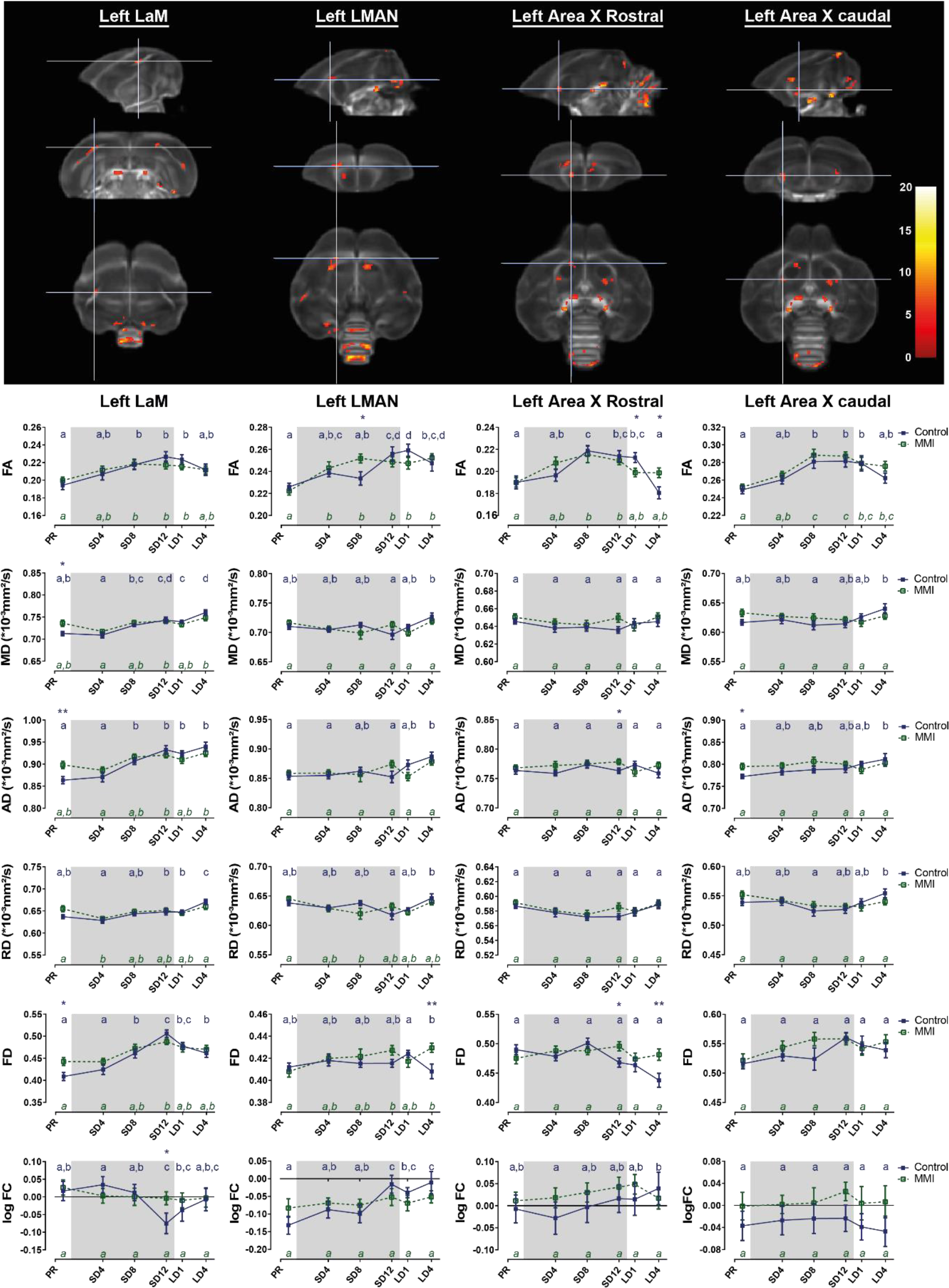
Summary of the significant longitudinal changes over time in FA, MD, AD, RD, FD and log FC extracted from ROI-based clusters at level of the left LaM, LMAN, Area X rostral and caudal surroundings.

**Figure 7 – figure supplement 1:**
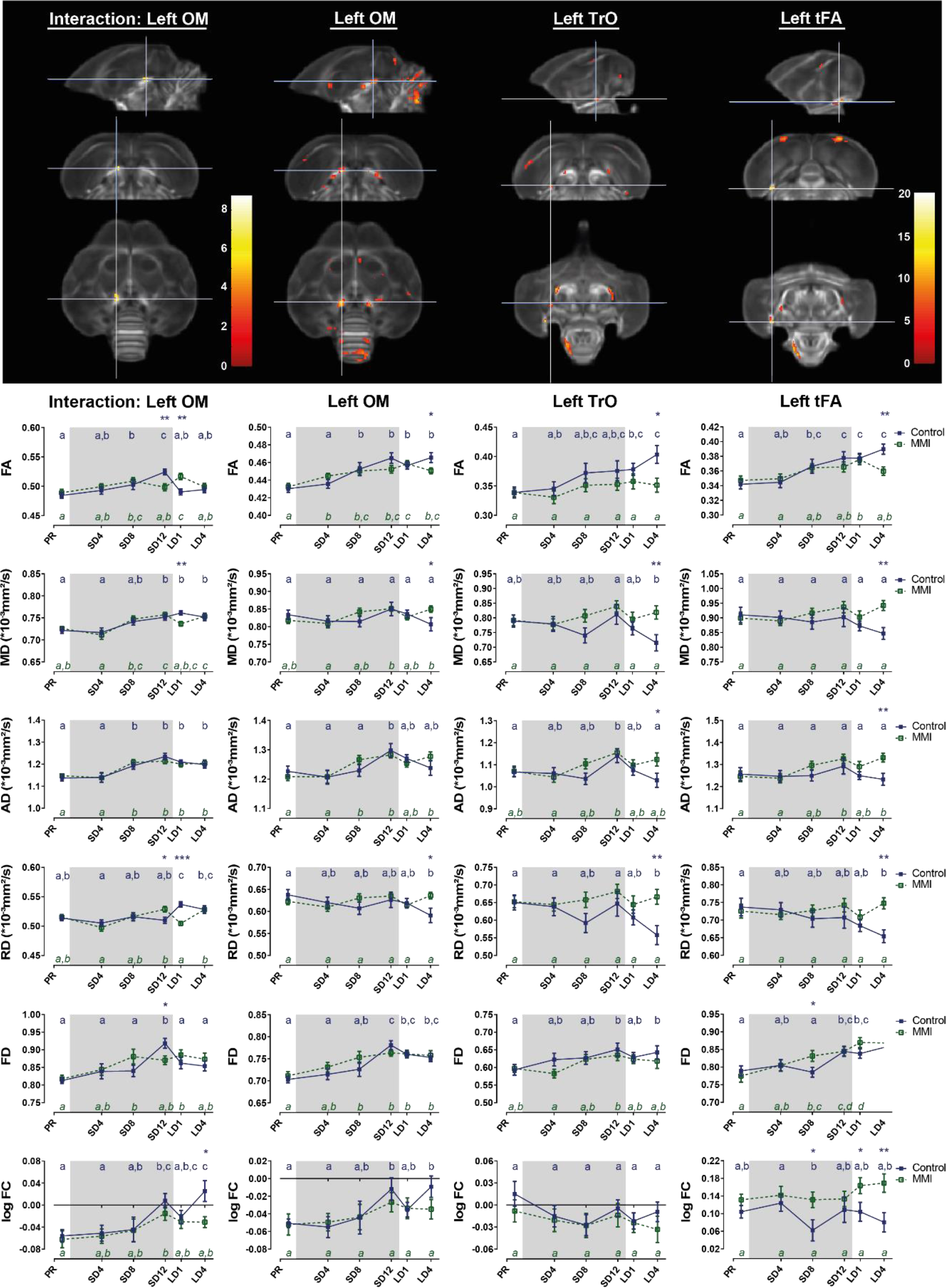
Summary of the significant longitudinal changes over time in FA, MD, AD, RD, FD and log FC extracted from ROI-based clusters at level of the left OM, TrO and tFA.

**Figure 7 – figure supplement 1:**
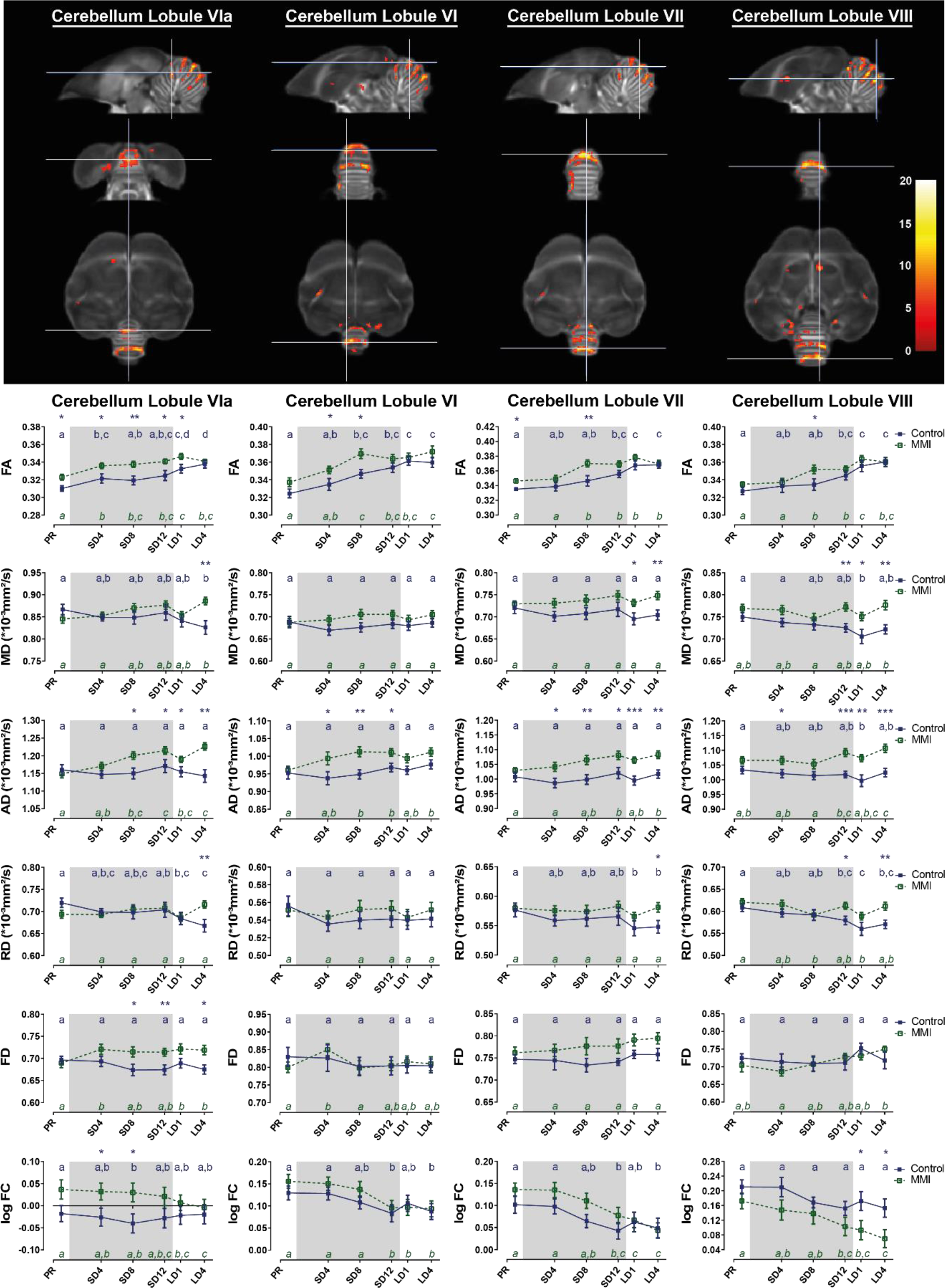
Summary of the significant longitudinal changes over time in FA, MD, AD, RD, FD and log FC extracted from ROI-based clusters at level of the cerebellar lobules VIa, VI, VII and VIII.

